# Non-plastic gene expression underlies root phenotypes involved in drought adaptation in *Vitis spp.*

**DOI:** 10.64898/2026.07.09.737455

**Authors:** Chedid Elsa, Patin R. Etienne, Tran Joseph, Marina de Miguel

**Affiliations:** EGFV, Univ. Bordeaux, Bordeaux Sciences Agro, INRAE, ISVV, F-33882, Villenave d’Ornon, France

**Keywords:** constitutive, drought, osmotic adjustment, plasticity, secondary metabolism, transcriptomic

## Abstract

Drought is a major abiotic stress threatening plant productivity and agricultural sustainability, yet the molecular mechanisms underlying adaptive root responses to water deficit in the water use strategies continuum remain insufficiently understood, particularly in perennial crops.

In this study, we explored drought responses in nine accessions belonging to three wild *Vitis* species (*V. acerifolia*, *V. candicans*, and *V. doaniana*) displaying variying drought-response strategies. Plants were subjected to moderate drought stress (40% soil water content) for three weeks under greenhouse conditions. By integrating physiological, metabolic, and transcriptomic analyses, we aimed to identify both conserved and species-specific mechanisms associated with drought adaptation.

Differential expression analyses revealed a conserved core set of drought-responsive genes shared among species, including genes involved in abscisic acid signaling, reactive oxygen species detoxification, solute transport, and plant defense. In parallel, each species exhibited distinct transcriptional and metabolic signatures reflecting alternative adaptive strategies related to osmoregulation, and oxidative stress mitigation. Weighted gene co-expression network analysis (WGCNA) further revealed significant associations between constitutive, non-plastic gene expression and root phenotypic traits.

Overall, our findings demonstrate that wild *Vitis* species rely on both conserved stress-responsive pathways and species-specific constitutive regulation to cope with drought stress. These results highlight the importance of root-associated traits and intrinsic regulatory networks in shaping drought adaptation and provide new targets for the development of drought-resilient grapevine rootstocks.

## Introduction

Drought is a major constraint on plant growth, productivity, and survival, with increasing impacts under ongoing climate change (IPCC, 2023). As drought events become more frequent and severe, understanding the mechanisms that underpin plant responses to water limitation has become a central objective in plant biology (Codon et al. 2004; Lesk et al. 2016). While aboveground symptoms of drought are readily observable, roots constitute the primary interface with the soil environment and act as the first sensors of declining water availability. Through their roles in water uptake, hydraulic regulation, and long-distance signaling, roots are key drivers of whole-plant responses to drought (Uga et al., 2013; Comas et al., 2013). Yet, compared with shoots, root-specific mechanisms remain comparatively understudied, leaving major gaps in knowledge regarding how roots perceive, integrate, and respond to water limitation, especially in perennial crops.

Drought responses are often characterized by dynamic physiological and transcriptional changes (Shinozaki & Yamaguchi-Shinozaki, 2007). Core mechanisms—including abscisic acid (ABA) signaling, reactive oxygen species (ROS) homeostasis, osmotic adjustment, and cell wall remodeling—are widely conserved across species (Carvalho et al., 2015; Chaudhry and Sidhu, 2022). These responses are typically interpreted as transcriptionally plastic, enabling plants to dynamically adjust to fluctuating environments (Konecny et al., 2024; Lin et al., 2023). However, increasing evidence suggests that constitutive genotype-dependent gene expression may also play a major role in shaping drought tolerance, independently of environmental cues (de Maria et al. 2020, Steele et al. 2026). The relative contribution of plastic versus non-plastic components remains poorly understood. Moreover, results obtained from above-ground organs may not be directly comparable to those from roots, which develop in substantially more heterogeneous environments. It is even possible that the molecular reaction may even be reversed under water stress (Gargallo-Garriga et al. 2014).

At the whole-plant level, drought adaptation is often described along a continuum between conservative and acquisitive water-use strategies. Conservative genotypes limit water loss through rapid stomatal closure (Dayer et al., 2017; Tardieu & Simonneau, 1998), reduced hydraulic conductivity (Coupel-Ledru et al., 2017, Pou et al., 2012), and enhanced stress-protective mechanisms, such us osmotic adjustment and lignin biosynthesis (Hochberg et al., 2013; Hochberg et al., 2018). On the other side, acquisitive genotypes maintain gas exchange and growth for longer periods under water deficit, at the cost of increased vulnerability to severe stress (Hochberg et al., 2018). These contrasting strategies reflect trade-offs between growth and survival (Grime, 1974). However, the extent to which such strategies are determined by transcriptional plasticity versus intrinsic, genotype-dependent gene expression remains poorly understood.

Grapevine wild relatives (*Vitis* spp.) provides a powerful system to address these questions because they are considered closely related taxa that share much of their evolutionary history but differ in ecological preferences and stress tolerance (Cantu et al. 2024). This allows to capture natural variation across environments while avoiding phylogenetic noise in the identification of genetic and regulatory mechanisms underlying adaptive traits. In addition, grapevine wild relatives are a treasure for viticulture (Atak, 2025). In cultivated systems, grapevines are grafted onto North American wild Vitis accessions or F1 hybrids naturally resistant to Phylloxera (Chen et al. 2024). Rootstocks play a critical role in modulating water uptake (Gambetta et al. 2013), hydraulic conductance (Flor et al. 2025), and stress signaling (Cookson et al. 2013 ; Gautier et al. 2020,2021). Despite their importance, most commercial rootstocks derive from a limited set of North American species, constraining the genetic diversity available for improving drought tolerance (Riaz et al. 2019). The evolutionary diversity, combined with their genetic relatedness, makes wild *Vitis* spp. particularly suitable for comparative transcriptomics aimed at dissecting drought-response mechanisms and identifying targets for grapevine breeding.

Previous transcriptomic studies in grapevine have identified key drought-responsive pathways, particularly involving ABA signaling, aquaporins, and secondary metabolism (Khadka et al., 2019, Cochetel et al. 2020, Hanzouli et al. 2025, Rossdeutsch et al. 2016). However, these studies have largely focused on cultivated genotypes or single species and have mainly emphasized stress-induced responses. Inter-specific comparative transcriptomics remain scarce, especially for root tissues. As a result, it is still unclear how conserved drought-responsive pathways interact with constitutive and plastic species-specific regulatory mechanisms, and how these differences translate into contrasting phenotypic strategies.

In this study, we used a comparative, multi-omics approach to investigate root drought responses across three wild *Vitis* species. Rather than contrasting extreme genotypes, we focused on subtle differences among species to capture the diversity of the water use strategies continuum. We tested the hypothesis that drought adaptation relies not only on conserved, plastic transcriptional responses but also on species-specific, non-plastic gene expression patterns that shape root morphology and physiology related to drought tolerance. Specifically, we aimed to (i) identify a conserved set of drought-responsive genes in roots, (ii) characterize species-specific transcriptional pathways, and (iii) link transcriptomic root responses to root and shoot phenotypes. By integrating transcriptomic, metabolic, and physiological data, this work provides new insights into the balance between plasticity and intrinsic regulation in plant drought adaptation and highlights the role of constitutive traits as targets for improving resilience.

## Material & Methods

### Plant material and drought experiment

Three wild Vitis *species* with different behavior under water stress were selected according to Patin et al. (2025). Each species was represented by three accessions. All the accessions were chosen from European germplasm collections and propagated as described in Supplementary Table 1.

The drought experiment was described in Patin et al. (2025). Briefly, the experiment was conducted with 6 months old cuttings in the Bordo greenhouse (INRAE EGFV). Six replicates per accession were randomly divided into six blocks. Three blocks were assigned to the well-watered treatment (WW) and 3 blocks to water deficit treatment (WD). As a result, there was one replicate per accession per block. Plants in the WD treatment were initially watered to full capacity and then water availability was progressively decreased to reach a target soil water content (SWC) of 40%. Throughout the experiments, the amount of water in the soil was determined by weighing the pot daily on balances (CH15R11, OHAUS type CHAMP, Nänikon, Switzerland, precision 1 g) (Sadok et al., 2007). WD pots were irrigated automatically mid-morning, to achieve the target water content. Plants were maintained under 40% SWC for 15 days. Plants in the WW treatment were manually irrigated four times per week to meet not limiting conditions. Harvest was done during 5 consecutive days in order to allow all plants to spend 15 days under the fixed SWC. Every WD replicate per accession was harvested at the same time than a replicate of the same accession from the WW treatment.

### Root and shoot phenotypes

Stomatal conductance to water vapor (gs, mol/m2/s) was measured in all the plants once per week. We measured three blocks per day, thus measuring all the plants lasted 2 days. We estimated Electron Transport Rate (ETR, µmol/m2/s) and Quantum Efficiency of Photosystem II in light conditions (⎞PSII) through chlorophyll fluorescence measurements. gs and chlorophyll fluorescence parameters were measured with a porometer/fluorometer (LICOR LI600, Lincoln, NE, USA) in a time slot between 10 a.m. and 12 p.m (at local time) on a fully developed leaf. During the third week of the experiment ETR and φPSII were measured in the three blocks of the WD treatment, but only on one block of the WW treatment. Having only one replicate per genotype on the WW treatment prevented significant differences between treatments for these traits in the third week of the experiment from being estimated. Consequently, shoot traits measured on the third week of the experiment were presented in Figure 1 but were not included in subsequent co-expresion analysis (see WGCNA subsection).

**Figure 1:**
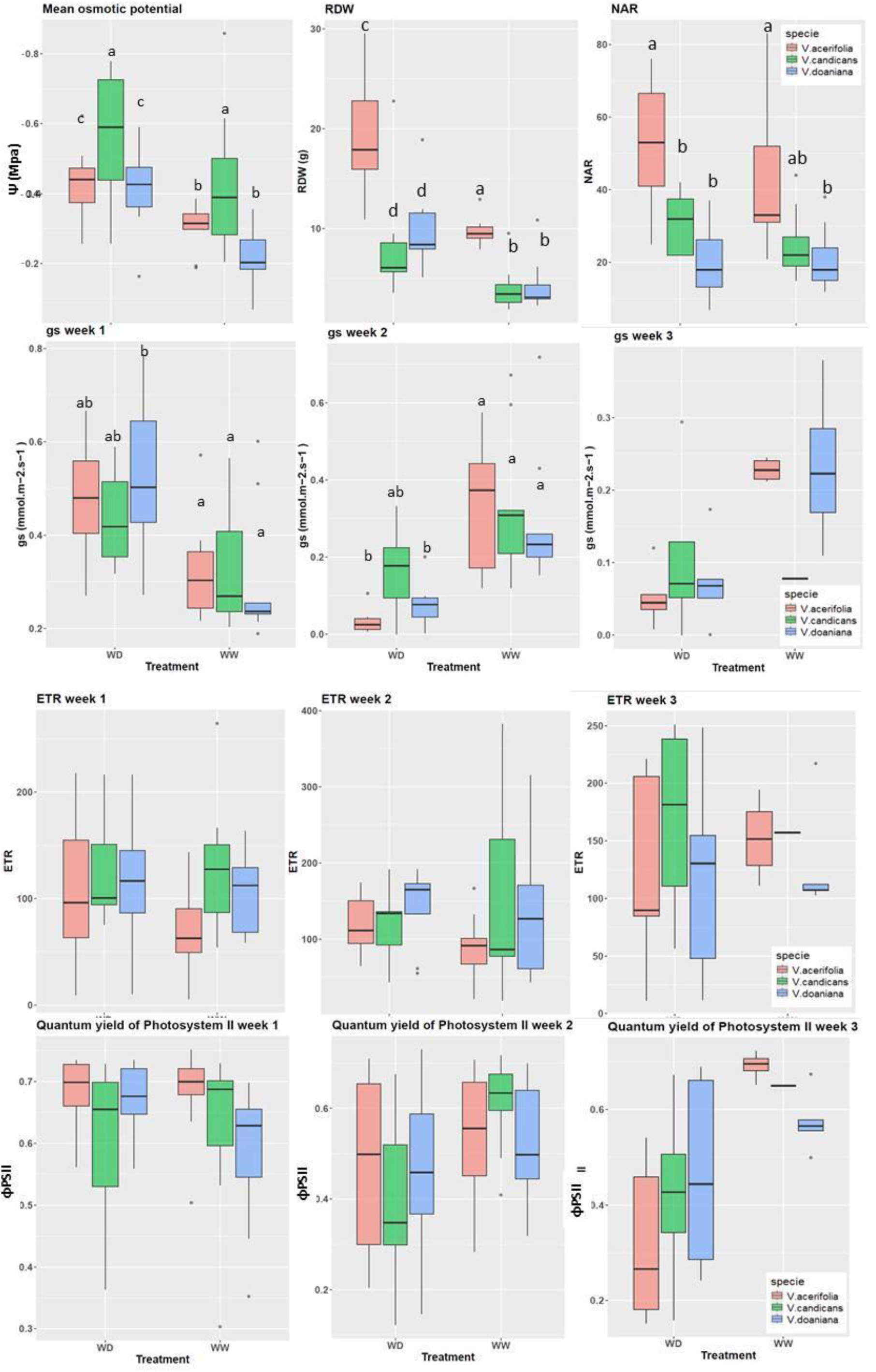
Boxplots of phenotypic traits for three Vitis species under well-watered (WW) and water deficit (WD) conditions over the course of a three-week experiment. V. acerifolia is pink, V. candicans is green, and V. doaniana is blue. ETR and and ⎞ PSII of week 3 were measured in three replicates for the WD treatment, but only one for the WW treatment. Having only one replicate per genotype on the WW treatment prevented significant differences between treatments from being estimated.

At the end of the experiment roots were harvested and the number of adventitious roots (NAR) was counted. We estimated root biomass as the root dry weight (RDW) after the root was oven dried at 80°C for 48 hours. Root osmotic potential at full-turgor was estimated with a Osmomat 030 osmometer at the root tip (Gonotec GMBH, Berlin, Germany) as described in Patin et al. (2025).

Root polyphenols quantification was done according to Patin et al. 2025. A calibration curve was built with polyphenol standards, including 6 flavanols (Catechin, epicatechin, procyanidin B1, procyanidin B2, epigallocatechin, gallocatechin), one phenolic acid (salycilic acid), 1 flavonol (quercetin 3-glucoside), and 9 stilbenes (trans-resveratrol, trans-piceid, ε-viniferin, miyabenol, hopeaphenol, isohopeaphenol, pallidol, vitisin A, vitisin B). Additionally, forms of procyanidin B4, cis-resveratrol and cis-piceid were detected and quantified as Procyanidin B2, trans-resveratrol and trans-piceid equivalents, respectively.

### RNA extraction, cDNA library construction and Illumina sequencing

Total RNA from grapevine root samples of both WW and WD treatments were extracted using the Spectrum™ Plant Total RNA Kit (Sigma-Aldrich, Product No. STRN50). A total of 150 mg of material was extracted with 900 µL of lysis solution with 60 mg of polyvinylpolypyrrolidone. The resulting solution was extracted once with chloroform:isoamyl alcohol (24:1, v:v) and then RNA was extracted according to the manufacturer’s instructions. The cDNA libraries were constructed at Neurocentre Magendie-INSERM NGS platform at Bordeaux (France). Pair-end sequencing was done using an Illumina NextSeq 2000 P3 (San Diego, CA, USA) at the PGTB platform at Pierroton (France). Further, the raw reads were analysed using the nf-core/rnaseq v3.18.0 pipeline (Harshil et al. 2024). The sequence reads were aligned to the grapevine reference genome PN40024.V4 (Velt et al. 2020).

### Analysis of gene expression level and enrichment analysis

The count data were analyzed using R software (R Core Team, 2021). The counts were normalized using the trimmed mean of M values (TMM) method. Differential analysis was performed using the edgeR (Chen et al., 2025) R package. This package has been used to identify differentially expressed genes (DEGs) through a multifactor design method. Several contrasts were analyzed: i) a contrast of WW versus WD plants for each species studied to identify species-specific DEGs; ii) an ANOVA-like (Anodev) test to test for differential expression across multiple groups and identify DEGs with common behavior between the three species. For each contrast, the log₂ fold change, p-value and adjusted p-value were evaluated, and only transcripts with an adjusted p-value <0.05 were analyzed.

Gene Set Enrichment Analysis (GSEA) was performed to identify significantly enriched biological pathways associated with the differential gene expression profile. In GSEA, all genes are ranked using the fold-change metric and then checked to see if they belong to a specific set or pathway based on GO terms. GSEA was conducted using the GSEA function of the clusterProfiler package in the R software program with the default parameters. The enrichment results were considered significant if the FDR q-value was less than 0.05 and the nominal p-value was less than 0.05, and if there were a minimum of five genes per pathway. Visualization of enrichment plots was performed using the enrichplot package.

MapMan enrichment analysis was performed using the online Mercator4 (Schwacke et al. 2019). It identifies protein classes that are over- or under-represented within the full set of Mercator4 BINs. We used Two-sided Fisher’s exact test to identify significantly enriched protein categories with a 0.05 FDR-adjusted p-value cutoff.

KEGG (Kanehisa et al., 2007) was used to perform pathway enrichment analysis of DEGs and identify enriched pathways under WD. We used the ShinyGO 0.85.1 application (Ge et al., 2020).

### WGCNA

We did a co-expression analysis of the gene expression data, and phenotypic traits. The co-expression analysis was performed using the Weighted Gene Co-expression Network Analysis (WGCNA) package in R (Langfelder and Horvath, 2008) to obtain hierarchical clustering and identify co-expressed genes (hub genes). The pickSoftThreshold function was used to determine the optimal soft-thresholding power (Langfelder and Horvath, 2008). For each analysis, the lowest power with the scale-free topology fit index was used. Soft powers b = 8 were selected using the function pickSoftThreshold. The adjacency matrix was transformed into a topological overlap matrix (TOM) and the corresponding dissimilarity matrix (1-TOM). Afterwards, a hierarchical clustering dendrogram of the 1-TOM matrix was constructed to classify genes with similar expression levels into different gene co-expression modules. Modules were merged by using a criterion of MEDissThres = 0.25. Finally, the relationships between each module and phenotypic traits that were significantly different among sites for each genotype were estimated by Pearson’s correlation using the module eigengene values. Modules with high correlation coefficients and a correlation adjusted p-value ≤ 0.05 were selected for subsequent analysis.

Shoot traits measured on the third week of the experiment were not included in WGCNA analysis because they were measured only in one replicate of the WW treatment during the last week of the experiment.

### Random Forest Analysis

To classify species based on gene expression profiles, we employed a random forest (RF) classification approach (Breiman L., 2001) using R’s randomForest and caret packages (Kuhn, 2008; Liaw & Wiener, 2002). We used the normalized gene expression matrix and genes with low variance across samples were removed to reduce noise. The RF model was trained on part of the data set, using 500 trees and default parameters. Model performance was evaluated using 5-fold cross-validation, and classification accuracy, confusion matrices, and variable importance scores were computed to assess model robustness and identify key genes contributing to species discrimination. The RF calculation was repeated 100 times to ensure the accuracy of the model and finally mean importance of all genes was calculated. All analyses were conducted in R v4.3.0.

## Results

Illumina sequencing was performed to identify changes in gene expression in the roots of three grapevine species following a period of water deficit. A total of 21,628 genes were retained for DEG analysis. Genes with a fold change of at least 2 and a False Discovery Rate (FDR) of less than 0.05 were defined as differentially expressed genes.

Principal component analysis (PCA) was performed to identify the main source of variation in the dataset (Supplementary Figure S1). The PCA plot based on normalized gene expression data showed separation of the samples along the first two principal components, which explained 18% and 14% of the variance, respectively. Ellipses enclose samples of the same species and treatment, indicating variance within each factor combination. Initially, samples from WW and WD were grouped into different clusters, reflecting the impact of the WD treatment on gene expression. The WD treatment showed greater dispersion than the WW groups, indicating higher variation in gene expression among genotypes under WD. *V. candicans* under WD showed the highest level of intraspecific variation. Overall, this PCA plot demonstrates that differences in gene expression were observed between watering treatments and among different species.

### Phenotypic response to drought

During the first week of the experiment, SWC decreased progressively, with most plants reaching the 40% SWC threshold after 10 days. Growth rates were maintained during the first week, but decreased significantly during the second week and ceased completely by the end of the experiment (Patin et al., 2025). Transpiration rates decreased throughout the experiment for all species (Patin et al., 2025).

The *V. acerifolia* and *V. doaniana* plants showed relatively similar responses to the water deficit. The mean osmotic potential (Ψ) decreased significantly in the roots of both species under WD (Figure 1). Both species also showed a significant decrease in stomatal conductance to water vapor (g_s_) during the second week of the WD period, with a drop from 0.3 to 0.08 mmol m⁻².s⁻¹ in *V. doaniana* and from 0.33 to 0.03 mmol m⁻².s⁻¹ in *V. acerifolia* (Figure 1). gs was higher in WD than in WW during the first week of the experiment because the WW plants were closer to the greenhouse fan. The root dry weight (RDW) of both species was higher under WD, while the number of adventitious roots (NAR) did not change under WD. However, *V. acerifolia* had significantly higher NAR and RDW than the other species in both treatments (Figure 1).

For *V. candicans*, high intra-specific variance prevented significant differences in ψs from being detected among treatments. However, the overall mean by treatment (−0.43 MPa in WW and -0.56 MPa in WD) was significantly lower than for *V. acerifolia* and *V. doaniana*. Additionally, no significant decrease in g_s_ in response to WD was observed for this species. RDW increased under WD, while NAR remained unchanged.

Although there was a slight decrease in the quantum yield of photosystem II (ϕPSII) under WD, no significant differences were observed between species or treatments during the first two weeks of stress. Similarly, the electron transport rate (ETR) did not change according to species or treatment, suggesting that the photochemical machinery was not impacted by the water deficit until the third week of stress.

A total of 20 polyphenols were quantified in the roots of the species studied under WW and WD conditions. A significant increase in the concentration of ε-viniferin, a resveratrol dimer, was observed in the three species under WD. This metabolite is considered a stress marker (Langcake & Pryce, 1981).

*V. candicans* was the species that accumulated polyphenols more frequently under WD. Indeed, a significant increase in the quantity of t-piceid, catechin, epicatechin, vitisin A and B, and procyanidins B1, B2 and B4 was observed in *V. candicans*, whereas the quantities remained constant in the other species, except for trans-piceid, which also increased in *V. acerifolia* (Figure 2).

**Figure 2:**
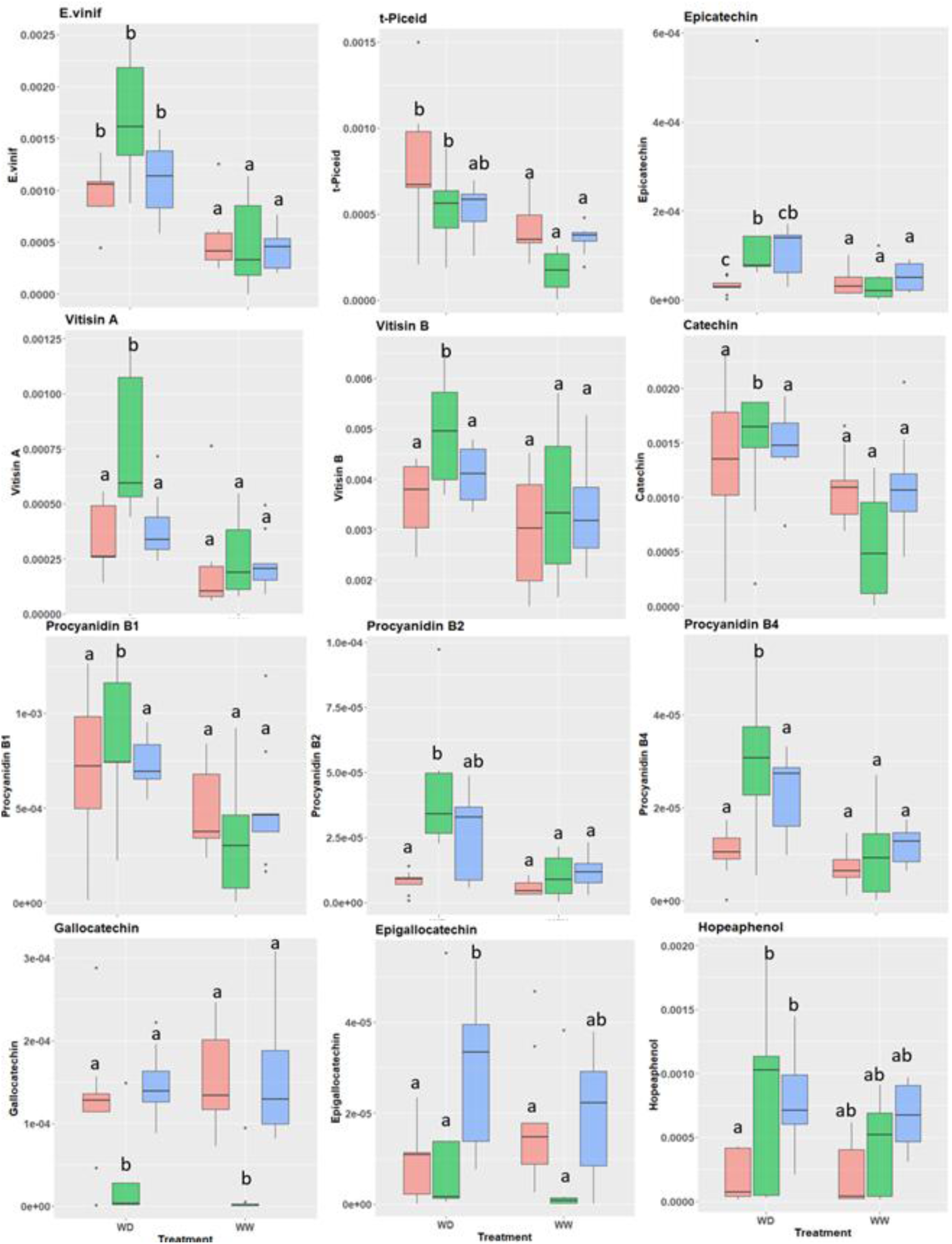
Boxplots of polyphenols quantified in roots for three Vitis species under well-watered (WW) and water deficit (WD) conditions at the end of a three-week experiment. V. acerifolia is pink, V. candicans is green, and V. doaniana is blue.

The quantities of gallocatechin and epigallocatechin varied significantly according to species (Figure 2). *V. candicans* showed lower quantities of gallocatechin than the other species in both conditions. The quantity of epigallocatechin in the roots of *V. doaniana* differed significantly from that in *V. acerifolia* and *V. candicans* under WD.

No differences were observed in salicylic acid, isohopaneol, pallidol, quercetin 3-O-glucoside, resveratrol and trans-piceid between species or treatments (Supplementary Figure S3).

### Conserved response to drought in gene expression across *Vitis* spp

The ANOVA-like differential analysis was conducted to identify the common DEGs between the three species under WD. This analysis allowed to identify genes with the same behavior among the species. In total, 2507 DEGs showed this trend, with a clear effect of water deficit on gene expression (Figure 3). The most up-regulated gene coded for glutaredoxin-C6 which is a key player in oxidative stress response and cellular redox homeostasis (Shutian Li, 2014; Wu et al. 2017). The most down-regulated gene coded for STY17, a Serine/threonine-protein kinase, playing a crucial role in the phosphorylation of chloroplast precursor proteins (Martin et al. 2006; Lamberti et al. 2011; Eisa et al. 2019).

**Figure 3:**
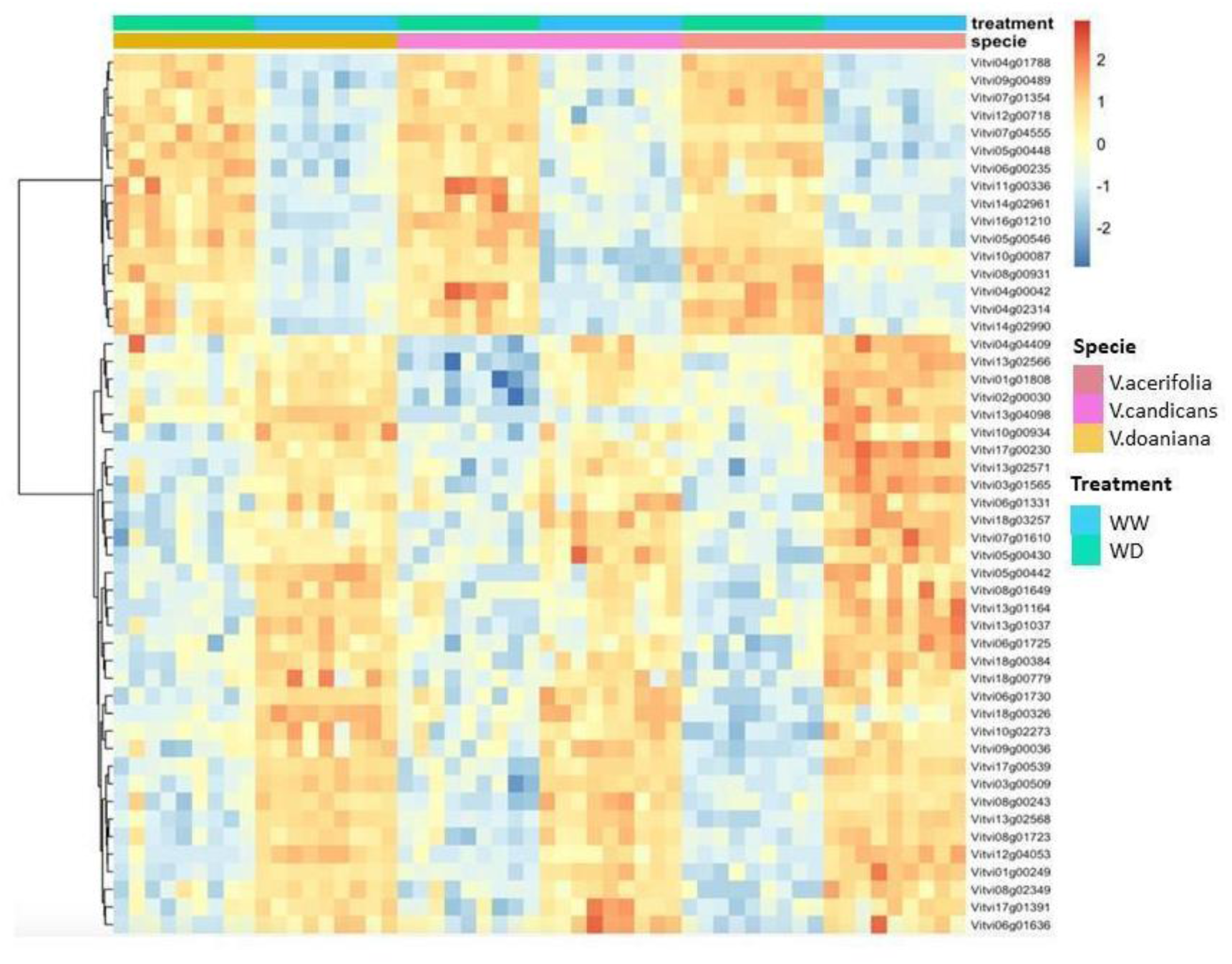
Heatmap for top 50 differentially expressed genes in the roots of three Vitis spp. under water deficit.

According to GSEA, the most enriched category of up-regulated DEGs was protein folding (Figure 4a). The protein folding enriched term, involved in protein homeostasis, included mainly small Hsp holdase chaperones (24 DEGs) (Supplementary Table 2).

**Figure 4:**
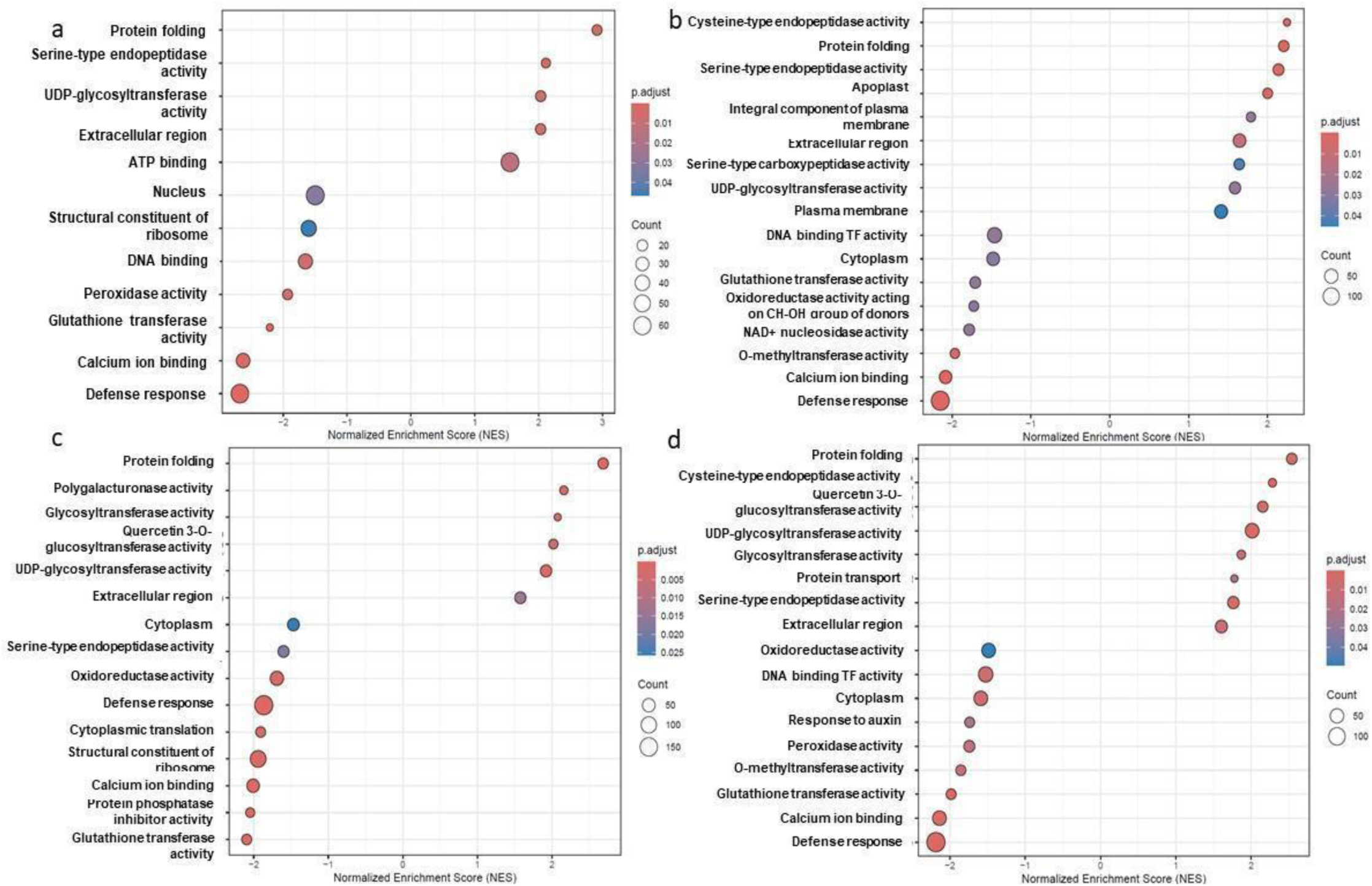
GO enrichment analysis of DEGs (a) conserved in the roots of three Vitis species, (b) in V.acerifolia, (c) in V. candicans and (d)in V. doaniana.

The highest number of upregulated genes were involved in ATP-binding (Figure 4a). The ATP-binding genes in the conserved response included mainly ABC solute transporters and transferase enzymes involved in protein phosphorylation and phytohormone signaling. These enzymes included TKL protein kinase family such as LRR receptor-like serine/threonine-protein kinase and CAMK protein kinase family. The Heat shock proteins (Hsp70 & Hsp90) were also part of the upregulated ATP-binding proteins. Belonging to the ATP-binding enriched term, the MFS superfamily was the most represented carrier-mediated transporters (45 DEGs) and it included several important families such as Sugars Will Eventually be Exported Transporters (7 SWEET) and Nitrate Transporter 1/Peptide Transporter (15 NRT1/PTR) (Supplementary Table 2).

The GSEA revealed the downregulation of genes belonging to several GO terms (Figure 4a) and the most downregulated genes belong to defense response and the Ca^2+^ signaling pathway (Figure 4a). The downregulated genes, belonging to DNA binding GO term, included several families of TF such as Ethylene-responsive (ERF) TF (50 DEGs), bHLH TF family (23 DEGs), MYB TF family (23 DEGs) and WRKY TF family (18 DEGs) (Supplementary Table 2).

MapMan enrichment analysis revealed that hormone metabolism, CHO metabolism, Phenylpropanoids metabolism, ABC and nitrate transport, ABA and ethylene signaling were enriched (Supplementary Table 3).

The KEGG enrichment analysis of the core response highlighted the enrichment of the flavonoid metabolism and biosynthesis of secondary metabolites along with starch, sucrose and nitrogen metabolism (Table 1). In this sense, we identified 59 DEGs from Phenylpropanoid Pathway (*e.g*. PAL, P450, 4CL, Isochorismate synthase, CCoAOMT and chalcone synthase) (Supplementary Table 4), which may play an important role as ROS scavengers. Interestingly, 37 DEGs regulating the photosynthetic activity were expressed in roots. They were mainly related to photophosphorylation (23 DEGs) and Calvin cycle (9 DEGs) (Supplementary File 2).

**Table 1:** KEGG enrichment analysis for DEGs in the core response and specific pathways of the three studied species (only enriched pathways unique to the species are shown). N Genes stand for Number of Genes enriched, Pathway Genes stands for total number of Genes in the Pathway, Fold Enrich stands for Fold Enrichment, FDR stands for False Discovery Rate.

|  | Pathway | N Genes | Pathway Genes | Fold Enrich | FDR |
| --- | --- | --- | --- | --- | --- |
| **Core response** | Nitrogen metabolism | 7 | 16 | 4.65 | 0.01 |
|  | Flavonoid biosynthesis | 7 | 26 | 3.36 | 0.05 |
|  | Biosynthesis of various plant secondary metabolites | 10 | 40 | 2.78 | 0.04 |
|  | Starch and sucrose metabolism | 19 | 72 | 2.56 | 0.01 |
|  | Plant-pathogen interaction | 25 | 128 | 1.98 | 0.01 |
|  | Biosynthesis of secondary metabolites | 101 | 701 | 1.38 | 0.01 |
| **V. acerifolia** | Metabolic pathways | 319 | 1239 | 1.18 | 0.01 |
|  | MAPK signaling pathway-plant | 30 | 87 | 1.83 | 0.01 |
|  | Plant hormone signal transduction | 58 | 200 | 1.53 | 0.01 |
| **V. candicans** | Ribosome | 68 | 166 | 3.36 | 5.31E-19 |
|  | Galactose metabolism | 10 | 26 | 3.31 | 0.01 |
|  | Glutathione metabolism | 18 | 66 | 2.43 | 0.01 |
| **V. doaniana** | Galactose metabolism | 11 | 26 | 3.94 | 0.00 |
|  | Pyruvate metabolism | 17 | 52 | 3.04 | 0.00 |
|  | Glycolysis/Gluconeogenesis | 21 | 72 | 2.73 | 0.00 |
|  | Carbon metabolism | 28 | 143 | 1.77 | 0.03 |
|  | Metabolic pathways | 182 | 1239 | 1.38 | 0.00 |

### Species-specific response

In addition to genes involved in the conserved response to WD among the three studied species, we identified the DEGs per species using the pairwise comparison (WW vs WD per species) and we focused on those DEGs only significant in each one of the three species (Supplementary Figure S3). We conducted a GSEA enrichment analysis on the differentially expressed genes per species to highlight the specific mechanisms that may be more pronounced in one species than the others.

#### V. acerifolia specific mechanisms

*V. acerifolia* was the species with the highest number of specific genes, with 2205 specific genes for a total of 5139 DEGs (Supplementary Figure S3). According to the GSEA, several processes were specifically relevant in *V. acerifolia* roots (Figure 4b). Transcriptional factor activity and the NAD+ nucleosidase activity downregulation were enriched exclusively in *V.acerifolia,* which may indicate a mechanism to save energy (Canto et al. 2015). The downregulation of the oxidoreductase and O-methyltransferase activities were also exclusively enriched in *V. acerifolia*.

The protein homeostasis was enriched in *V. acerifolia* with serine-type carboxypeptidase and cysteine-type endopeptidase categories being up-regulated. The cellular components mostly involved in *V. acerifolia* response were the plasma membrane, its integral component and the apoplast (Figure 3b). The genes belonging to the plasma membrane and its integral component were mostly solute transport genes. The genes involved in the apoplast in *V. acerifolia* were mainly involved in cell wall organization. For example, DIRIGENT Proteins that play crucial roles in cell wall reinforcement, stress response, and secondary metabolism especially in lignan and lignin biosynthesis were up-regulated, ESB1 (*Enhanced Suberin 1)* gene that plays a critical role in regulating suberin deposition in the Casparian strip (Hosmani et al. 2013) was also up-regulated.

In addition, we identified several genes specific to *V. acerifolia* that were not present in the list of conserved DEGs. Three aquaporin genes (Vitvi15g01110, Vitvi13g00255, Vitvi08g01038) in the solute transport category were specifically up regulated in *V. acerifolia* (PIP1-2, PIP2-5 and TIP). Also, a strong modulation of transcription factors was observed; 14 MYB related genes, 17 bHLH and 22 NAC were only differentiated in *V.acerifolia.* Regarding the subtilases, *V. acerifolia* also showed specific response with 19 specific genes out of 25 DEGs. Within the cell wall organization category, Vitvi01g01657 gene coding for a cuticular lipid transfer accessory factor (LTPG) was exclusively upregulated in *V. acerifolia*.

According to KEGG enrichment analysis, three pathways were exclusively enriched *V. acerifolia* (Table 1). They showed higher hormone and MAPK signalling pathways while the metabolic activity was lowered.

#### V. candicans specific mechanisms

For *V.candicans*, a total of 2720 DEGs were identified from which a set of 777 genes were species-specific (Supplementary Figure S3). Among these genes, 81 genes coding for ribosomal proteins were downregulated while their expressions did not change in the other two species. Also, 12 cytochrome P450 genes were found differentiated only in *V.candicans*.

Several significant GO enriched terms, specific to *V. candicans* were identified (Figure 4c). We observed an enrichment in the downregulation of genes related to cytoplasmic translation and ribosome function. This response could be related to an adjustment of the efficiency of translation, ribosome composition, and protein synthesis, that would enable cells to prioritize stress responses and conserve resources under water-limiting conditions (You et al. 2019; Son et al. 2023). The downregulation of protein phosphatase inhibitor activity involves the downregulation of ABA receptors.

On the other hand, several specific GO terms were up regulated; Glycosyltransferase activity and quercetin 3-O-glucosyltransferase activity. Finally, polygalacturonase (PG) activity upregulation was exclusively enriched in *V. candicans*. The PG genes are known to have an impact on root cell elongation through cell wall remodeling and pectin breakdown (Yang et al. 2018).

These results were also highlighted by the KEGG enrichment analysis (Table 1).

#### V. doaniana specific mechanisms

*V. doaniana* had a total of 2517 DEGs identified from which a set of 391 genes were species-specific (Supplementary Figure S3). The GO terms enrichment showed many processes in common with the other two species (Figure 4d). The enrichment in protein homeostasis (cysteine-type endopeptidase), transcription regulation and O-methyltransferase activities was in common with *V.acerifolia*. The enrichment in glycosyltranseferase and glucosyltransferase activities were in common with *V.candicans*.

The KEGG analysis showed an enrichment of Glycolysis and galactose pathways (Table 1) which can be upstream of pyruvate pathway. Also carbon metabolism was specifically enriched in *V. doaniana*.

Three processes were specifically enriched in the response to WD in *V. doaniana*: the peroxidase activity, auxin response and protein transport. The peroxidase and O-methyltransferase activities that normally mitigates oxidative stress (Caverzan et al. 2012; hafeez et al. 2021) were downregulated. Peroxidase activity also involves genes like lignin-forming anionic peroxidase that can regulate cell wall formation. So, the role of peroxidase in roots is not restricted to ROS scavenging but also to cell wall formation and growth regulation. Within the response to auxin we found downregulated SAUR (Small Auxin Up RNA) genes.

Early auxin-responsive proteins play roles in cell elongation and growth in response to auxin hormone (Ren et al. 2015). Finally, within the protein transport category, the upregulation of oligopeptide transporters (OPTs) can be highlighted because of their key role in nutrient uptake and stress adaptation (Lubkowitz M. 2011).

### Co-expression gene modules

Co-expression networks of weighted genes associated with the phenotypic traits were constructed by WGCNA based on 15000 genes. Those genes were selected based on the topmost variable genes using Median Absolute Deviation. They were grouped into 16 co-expressed modules (Table 2). Each set of highly correlated genes corresponded to a branch of the tree (Supplementary Figure S4). Module sizes ranged between 116 and 2777 genes (Table 2). In general, the connectivity in our modules was high but the heterogeneity was also high reflecting a mixture of hub and peripheral genes in each module (Table 2). The module centralization was low which also indicates the presence of several hub genes per module. These parameters indicate that the networks are complex and involve many key genes. The mean module membership was between 0.5 and 0.7.

**Table 2:**
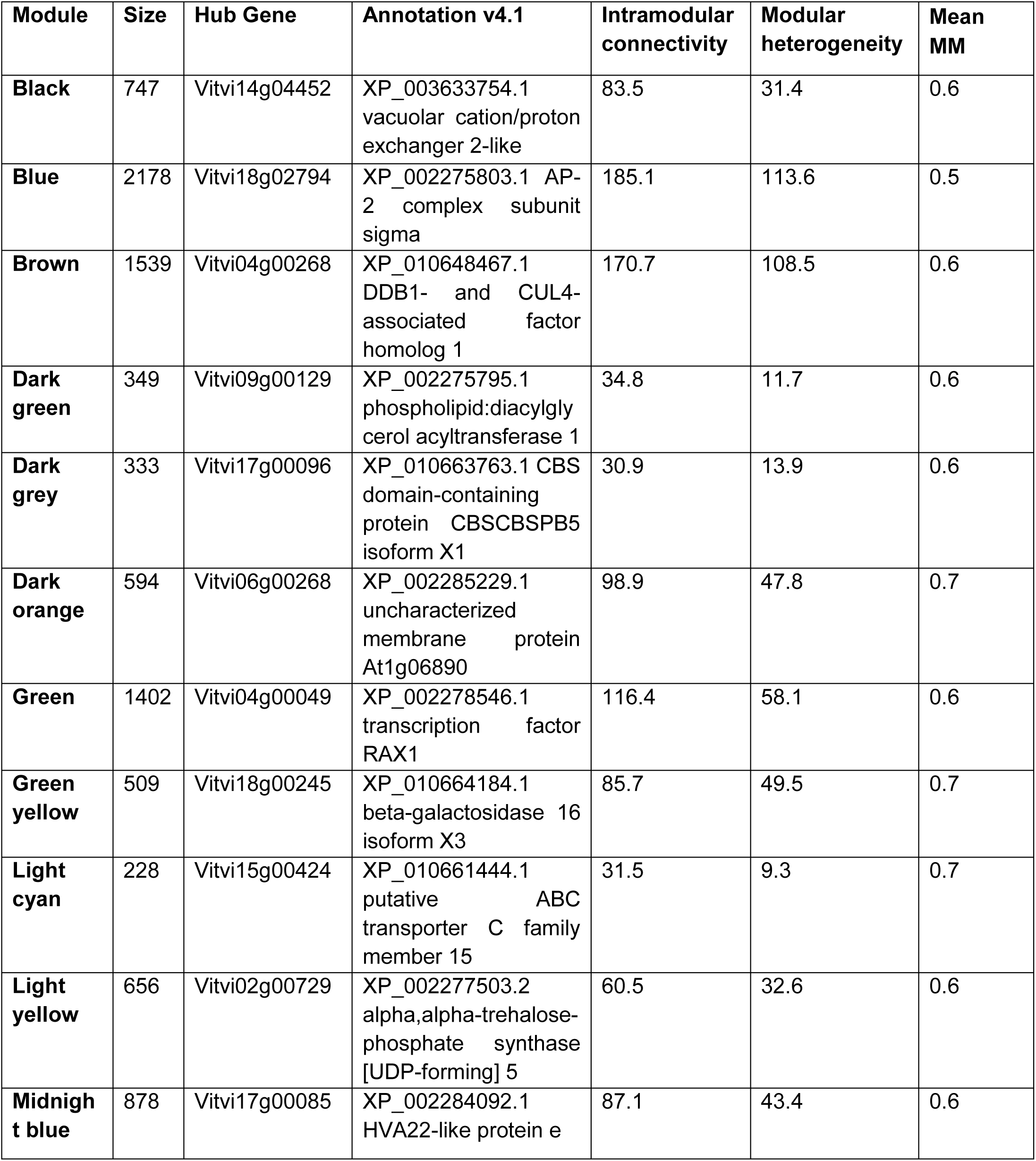
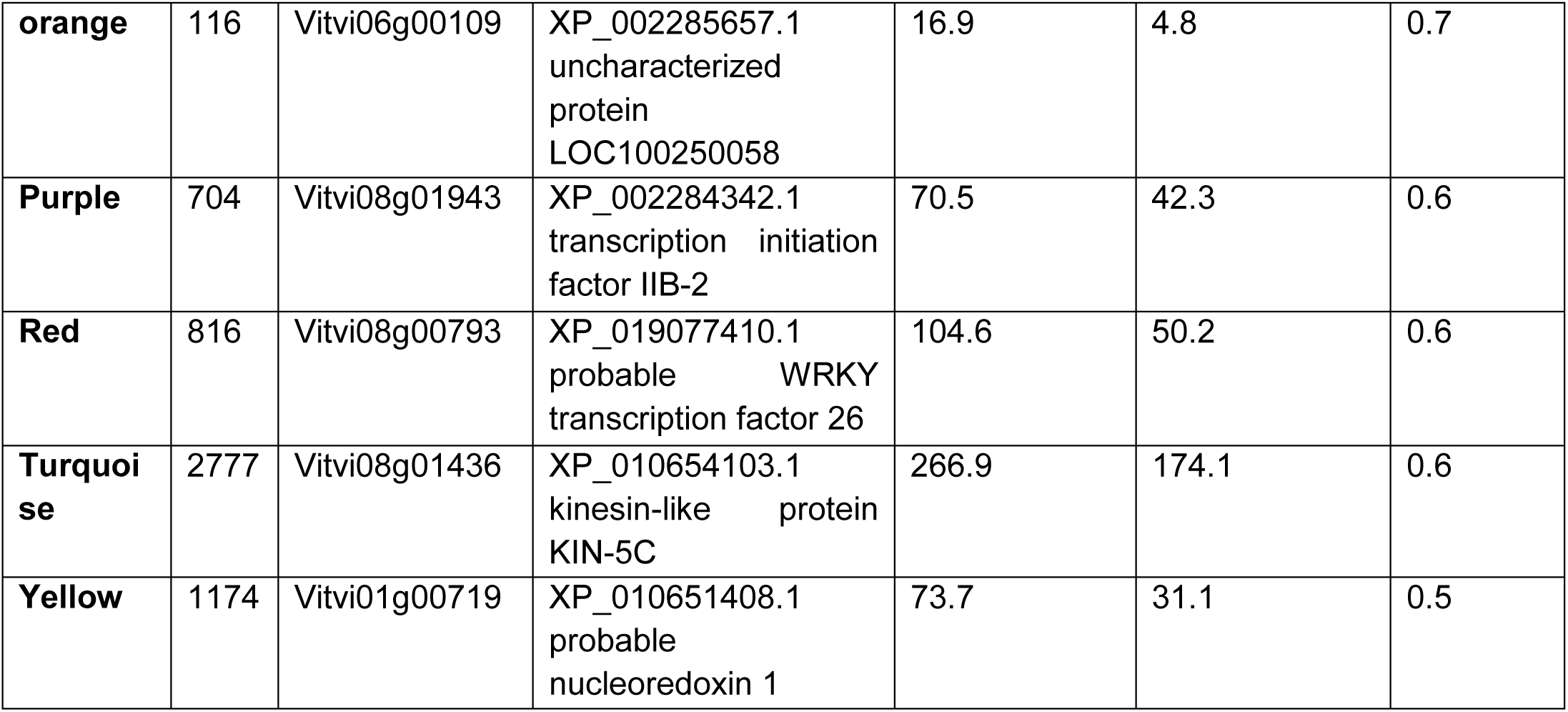
Module’s statistical parameters, hubgene identified and their annotations on PN40024.V4. MM is Module Membership.

According to module-trait correlations, we identified different significant correlations with root and shoot traits (Figure 5). Interestingly, we identified different module types: gene modules with expression profiles depending on water treatment and others where gene expression was species dependent (Figure 6).

**Figure 5:**
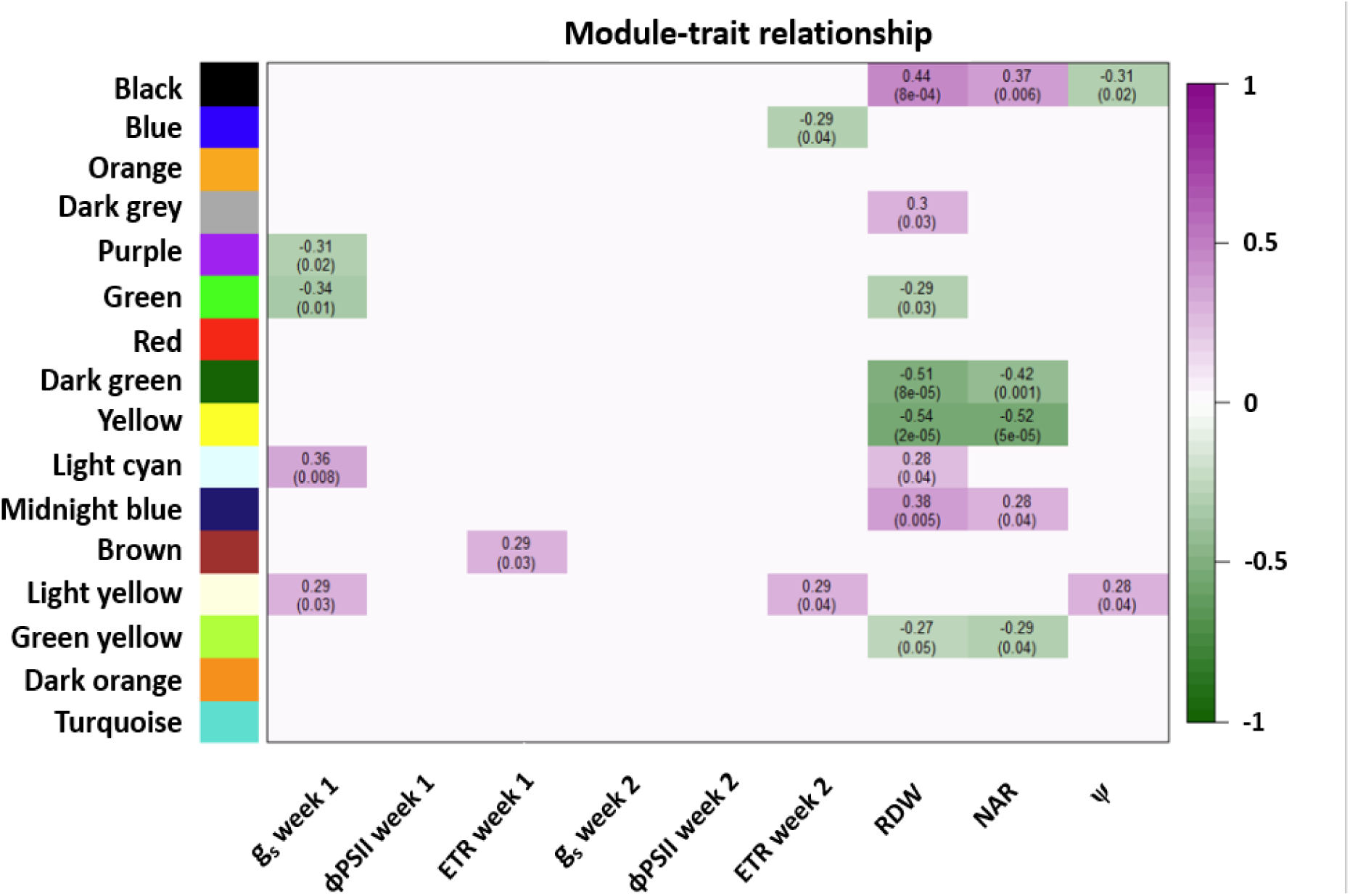
Module-trait correlations. Each column corresponds to a shoot or root phenotypic trait. Their association with module eigengenes (in rows) is represented by the Pearson correlation coefficient with the P-value within parentheses. The intensity of the cell color indicates the correlation coefficient: purple indicates a high positive correlation and green a high negative correlation.

**Figure 6:**
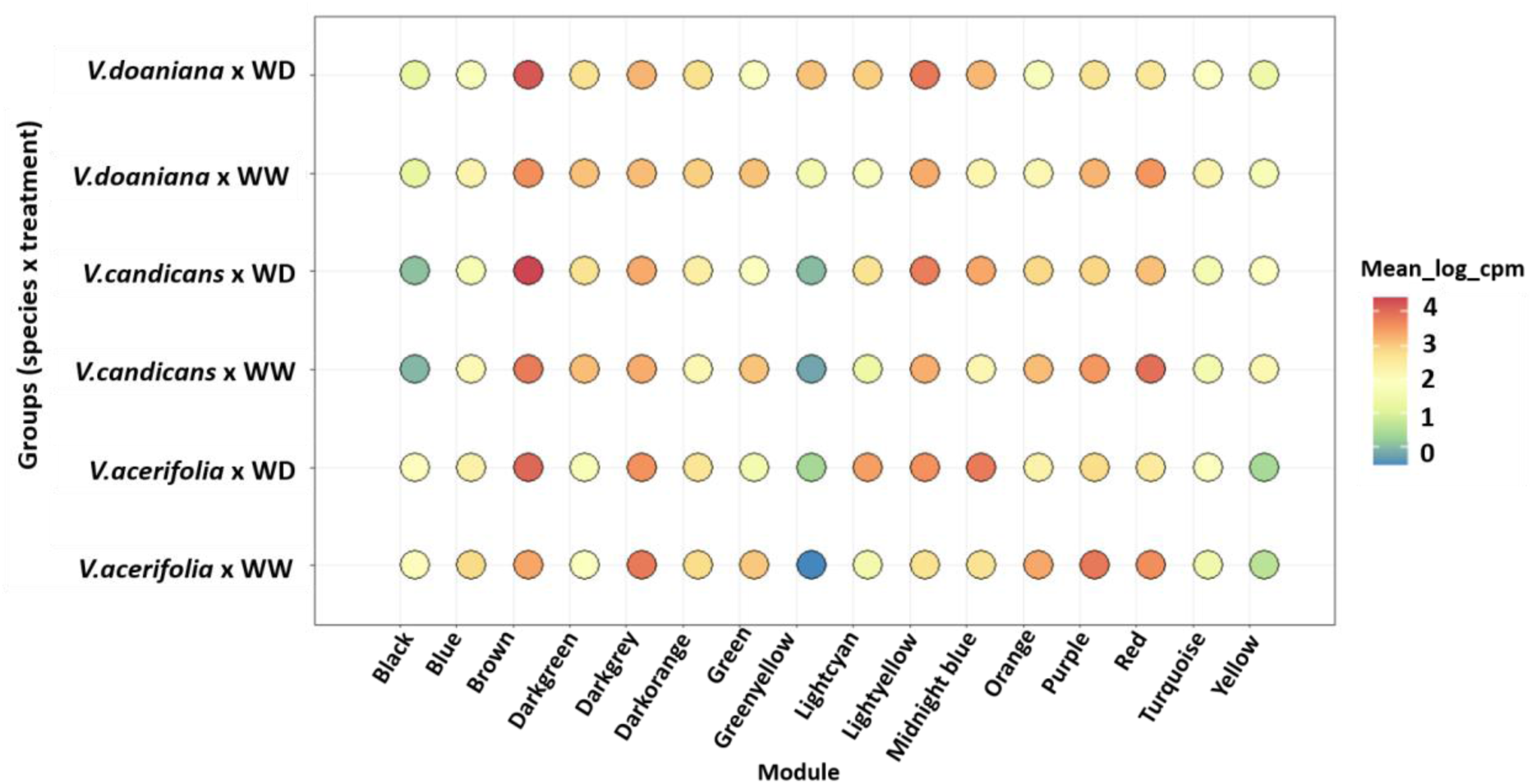
Expression level of each of the 16 modules in the 6 groups (3 species x 2 treatments) showing clusters that are species dependent or changed in response to WD.

**Figure 7:**
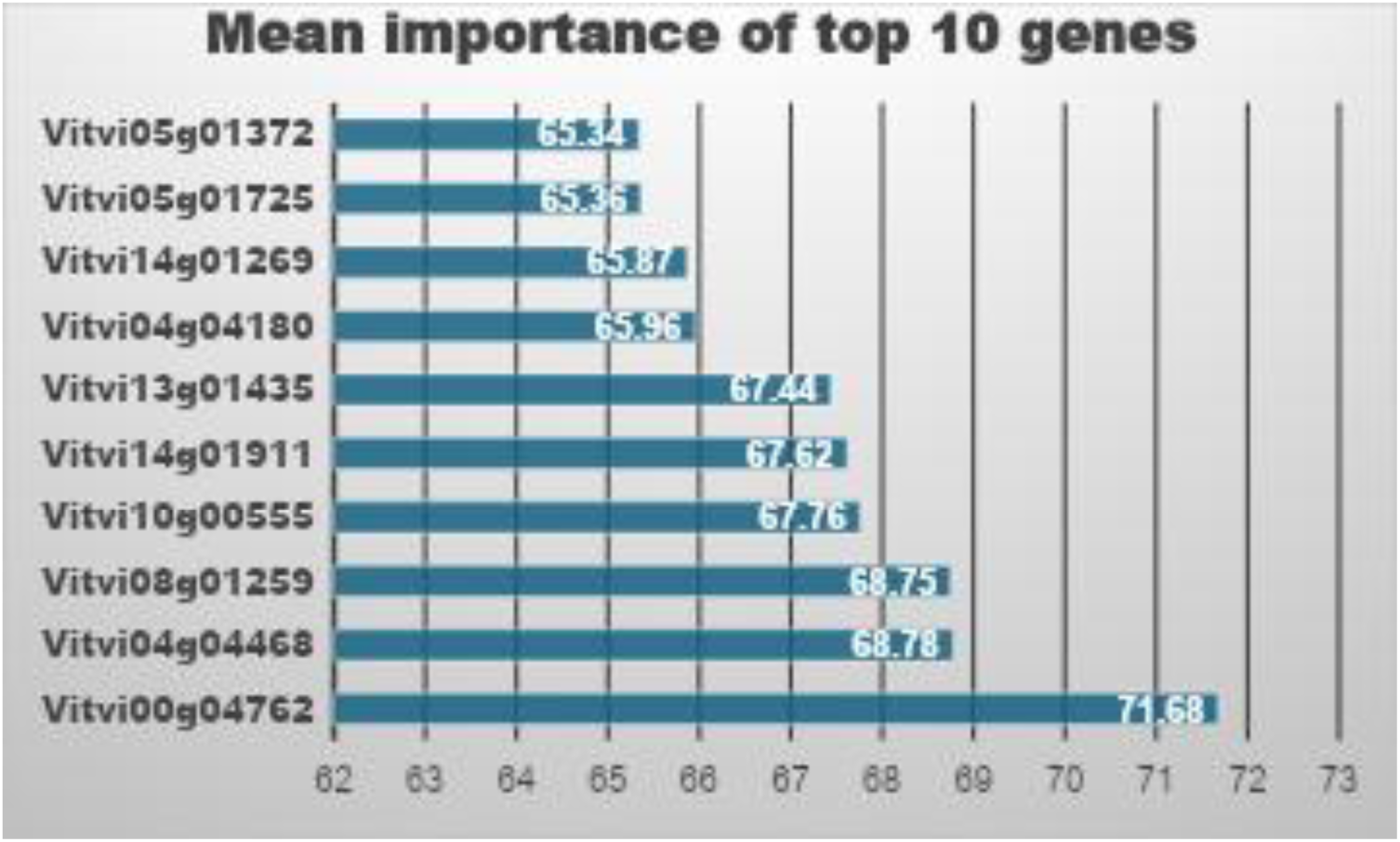
Top 10 genes mean importance to predict the species category (V. acerifolia, V. doaniana and V. candicans) calculated using Random Forest.

Three modules (light yellow, light cyan & green) were correlated with root and shoot traits and their expression depended mainly on the WD treatment (Figure 6). Those modules have the majority of the DEGs belonging to the conserved response among all three species. All the genes belonging to the light yellow module (i.e. 656 genes) were upregulated under WD in all three species. The most hub gene coded for a 2 alpha, alpha-trehalose-phosphate synthase (TPS), involved in the trehalose biosynthesis pathway, producing trehalose-6-phosphate, a key molecule in both metabolic regulation and stress protection (Tong-Ju Eh et al. 2024). This module correlated positively with ψ, ETR and g_s_ (Figure 5). The light cyan module had 220 out of 228 genes that were involved in the conserved response to WD among species. The genes had similar expression profiles, mostly up regulated under WD (Figure 6). The most hub gene coded for an ABC transporter. This module correlated positively with RDW and g_s_ (Figure 5) and nine polyphenols (Supplementary Figure S5). The Green module had 1042 genes out of 1420 also identified in the conserved response to WD. The genes in this module were down regulated under WD and their expressions was correlated negatively with RDW. The most hub gene coded for a transcription factor RAX1(REGULATOR OF AXILLARY MERISTEMS1) from the MYB TF family. Twelve other MYB TF were present in this module such as: RAX2, RAX3, MYB36, MYB4/4b and MYB1R1.

On the metabolic level, the light yellow and light cyan modules correlated positively with different procyanidins, flavonols and stilbenes (Supplementary Figure S5) reflecting the implication of those genes in the activation of secondary metabolism and antioxidant activity under WD. The downregulation of gene expression of the green module’s genes and its negative correlation with the same metabolites support this observation.

On the other hand, the gene modules dark grey, dark green, black and yellow have an expression profile dependent on the species rather than the water treatment. The black module had a positive correlation with RDW and NAR and negative correlation with ψ. The genes in black module had lower expression in *V. candicans* compared to *V. acerifolia* and *V. doaniana* (Figure 6). On the metabolic level, this module correlates with several metabolites from phenylpropanoid pathway. A strong positive correlation was detected with Gallocatechin and negative correlations with resveratrol and vitisins (Supplementary Figure S5). This may indicate that those genes are associated with the activation of Flavan-3-ol pathway rather than Stilbene pathway. Knowing that chalcone synthase (CHS) and stilbene synthase (STS) compete for common phenylpropanoid precursors, thereby influencing metabolic flux toward either flavonoid/procyanidin accumulation or stilbene production (Vannozzi et al. 2012), the lower expression of the genes channeling carbon into flavonoids in *V. candicans* compared to *V. acerifolia* and *V. doaniana* suggest two different adaptive defense strategies: stilbene phytoalexin defense, in *V. candicans*, vs. flavanol/procyanidin antioxidant defense, in the other species (Liu et al. 2021). The dark grey module had a positive correlation with RDW and its genes had slightly higher expression in *V.acerifolia*. This is a smaller module than the other ones, with a high number of unidentified genes. As a consequence, the enrichment analysis was not conclusive. The yellow and dark green modules had negative correlations with NAR and RDW. The genes of these modules had lower expression in *V. acerifolia* compared to the other species (Figure 6). On the metabolic level, the yellow and dark green modules showed positive correlations with several stilbene-associated metabolites, including trans-resveratrol, hopeaphenol, and viniferin-type compounds. Those genes were more expressed in *V. candicans* and *V. doaniana* than in *V. acerifolia* which suggest that these modules may be associated with inducible stilbene phytoalexin metabolism in *V. candicans* (Supplementary Figure S5), as also showed in the black module, (Flamini et al. 2013).

To link between the observed phenotypic differences among the species and gene expression, we searched in these modules for the genes with the highest positive or negative correlation coefficient with RDW, NAR and ψ. We found that the genes with highest correlation with RDW and NAR have significantly higher expression in *V. acerifolia* than *V. candicans* (Supplementary Table S5). These genes probably explain the higher root biomass in *V. acerifolia* (Figure 1). We found three TF (MADS-box TF PHERES 2), one methylation reader component (MBD9), one Nitrile-specifier protein, one EG45-like domain containing protein, one class I heat shock protein and one Subtilisin-like protease SBT5.3. The SBT5.3 has the larger expression difference between the species and it was found to have a role in early immune signaling (Figueiredo et al. 2016).

On the other hand, the genes with the highest negative correlation coefficient with ψ had higher expression in *V. acerifolia* and *V. doaniana* than *V. candicans* (Supplementary Table S5) which is in accordance with the differences in ψ at constitutive level (Figure 1). These genes coded for glycosyltranferases, CCoA-OMT, NCRP signaling, NRT1, Auxin related proteins.

### Gene expression variability among species

The RF model conducted on 15000 features matrices had an accuracy of 0.85. The genes mean importance (over 100 repetitions) of the top 10 genes ranged from 65.34 to 71.68% (Figure 7). These genes had very close values with less than 10% difference between the first and 20^th^ gene (Supplementary Table S6). The gene with the highest importance to differentiate the three *Vitis* species was Vitvi00g04762 that codes for a mitochondrial pyruvate carrier (MPC). The MPC controls the entry of pyruvate into mitochondria for the tricarboxylic acid (TCA) cycle and subsequent energy production (Monteiro-Batista et al. 2025). This gene with the second and the third discriminating genes, with 68% mean importance, (Vitvi04g04468 – not annotated; Vitvi08g01259 coding for oxidoreductase), were mostly expressed in *V. acerifolia*, less in *V. doaniana* and not expressed in *V. candicans*.

The genes with 67% mean importance (Vitvi10g00555, Vitvi14g01911 & Vitvi13g01435) had high expression in *V. candicans*, less in *V. doaniana* and very low expression in *V. acerifolia*.

The four genes with mean importance of 65% Vitvi04g04180 (GLR channel, Vitvi05g01372 (protein ROOT PRIMORDIUM DEFECTIVE 1) and Vitvi14g01269 (vacuolar cation/proton exchanger) were more expressed in *V. candicans* than the other species. Finally, Vitvi05g01725, coding for PHERES 2 TF, was highly expressed in *V. acerifolia*, less in *V. doaniana* and not at all in *V. candicans*.

Interestingly all these genes were part of the black and yellow gene modules. These modules, as we saw in the previous section, had non-plastic behavior to WD indicating they did not vary according to the environmental conditions.

## Discussion

Grapevine responses to drought are highly complex and are shaped by the interaction of multiple physiological traits, including stomatal regulation (Schultz, 2003), osmotic adjustment (Martorell et al., 2015), and root system architecture (Alsina et al., 2011), all of which contribute to the large variability observed among genotypes (Gambetta et al., 2020). Conventional transcriptomic studies of drought responses typically compare highly contrasting genotypes, usually categorized as drought-sensitive and drought-tolerant, to maximize differences in gene expression and identify pathways associated with extreme survival or growth responses. However, drought adaptation is better understood as a continuum of strategies rather than a binary response. In this study, we adopted a different approach by focusing on three wild *Vitis* species displaying contrasting, yet non-extreme, drought-response behaviors. This strategy allowed us to move beyond the identification of canonical drought-responsive pathways and examine how conserved plastic responses interact with species-specific constitutive adaptations. By focusing on the root transcriptomic response under a mid-term moderate water deficit (40% SWC), and integrating transcriptomic, metabolic, and physiological analyses, we identified a common drought-responsive signature associated with osmotic adjustment, oxidative stress mitigation, and metabolic reprogramming across species, while also revealing substantial differences in the coordination and intensity of these responses. Importantly, several pathways associated with root traits were already differentially regulated under well-watered conditions, suggesting that constitutive gene expression contributes to drought adaptation.

### Phenotypic response to drought

*Vitis acerifolia* and *V. doaniana* displayed similar physiological responses, characterized by an earlier reduction in stomatal conductance, and a more pronounced osmotic adjustment compared with *V. candicans*. In contrast, *V. candicans* exhibited greater intra-specific variability in osmotic adjustment and maintained stomatal opening for longer under water deficit conditions. Drought increased root biomass in all three species, although *V. acerifolia* maintained a significantly more developed root system overall. Photosynthetic activity remained relatively stable during the first two weeks of stress and declined only during the third week, suggesting that all species retained a certain degree of tolerance to moderate drought conditions (Chaves et al., 2010; Dayer et al., 2020). The drought-tolerance biomarker ε-viniferin (Hanzouli et al., 2024) accumulated in all three species, confirming that all genotypes experienced stress under our experimental conditions. However, polyphenol quantification revealed stronger metabolic activation in *V. candicans* relative to the other species, particularly *V. acerifolia*. This enhanced secondary metabolism may reflect a compensatory stress-response strategy associated with the maintenance of gas exchange under drought, potentially indicating a greater reliance on biochemical protection mechanisms to counterbalance oxidative damage.

A previous study conducted on the same genotypes analyzed here revealed that, under well-watered conditions, *V. candicans* exhibited a slower shoot growth rate than the other species, whereas drought-induced growth reduction became similar across all three species during the second and third weeks of stress (Patin et al., 2025). These results suggest that although carbon assimilation may be maintained for longer during short-term drought in *V. candicans*, this does not necessarily translate into sustained shoot growth. Our results further indicate that, under drought conditions, *V. candicans* preferentially allocated carbon toward the activation of phenylpropanoid metabolism and the accumulation of antioxidant compounds, especially stilbenes, which are known to contribute to oxidative stress mitigation. By contrast, *V. acerifolia* and *V. doaniana* appeared to rely more strongly on root osmotic adjustment, a strategy that may have supported greater root system development and potentially improved access to water from deeper soil layers (Alsina et al., 2011), although this hypothesis remains to be validated under field conditions. While the delayed stomatal closure observed in *V. candicans* may favor continued gas exchange during moderate stress, such a strategy can become disadvantageous under prolonged drought by accelerating water loss and increasing the risk of hydraulic failure and irreversible tissue damage (Delzon, 2015). Therefore, additional traits associated with hydraulic vulnerability and xylem safety margins should be investigated to better understand the adaptive significance of delayed stomatal regulation in this species.

Taken together, these results suggest that the three *Vitis* species occupy different positions along the water saving–water acquisitive continuum and they do not fit into discrete drought-response categories. *Vitis acerifolia* and *V. doaniana* displayed a strategy of belowground resource acquisition and water conservation through limiting transpirational water loss while maintaining water uptake capacity under declining soil moisture. In contrast, *V. candicans* drought responses suggest a comparatively more acquisitive strategy that prioritizes continued gas exchange and metabolic protection under moderate stress conditions. However, this strategy did not result in sustained shoot growth which suggests that the existence of trade-offs in carbon allocation modulates these continuum that should be interpreted multidimensionally.

### Core response to drought among *Vitis* species

Differential gene expression analysis enabled the identification of a conserved core gene set shared among the three *Vitis* species in response to drought stress in roots. Across all genotypes, drought induced the upregulation of genes associated with solute transport, protein folding, hormone signaling, and secondary metabolism. Among these responses, transporter-related gene families were particularly prominent, suggesting an active reconfiguration of membrane transport processes to maintain cellular homeostasis under water deficit. Notably, several ATP-binding cassette (ABC) transporter genes were strongly induced, consistent with reports in other plant species under drought stress (Yang et al., 2024; Yadav et al., 2025). In pear (*Pyrus bretschneideri*), numerous *PbrABC* genes were similarly upregulated following water deficit, where they were proposed to participate in the transport of signaling molecules, detoxification processes, and stress-related homeostasis (Kou et al., 2024). In grapevine, however, studies specifically addressing the role of ABC transporters in roots remain scarce.

The relevance of ABC transporters in the drought response was further supported by co-expression network analysis, where the light cyan module, enriched in transporter-related genes, showed significant correlations with stomatal conductance and root biomass. ABC transporters involved in abscisic acid (ABA) transport are known to mediate root-to-shoot signaling by facilitating ABA translocation from roots to aerial tissues (Borghi et al., 2015). This mechanism is consistent with findings in *Arabidopsis*, where ABCG-type transporters regulate stomatal responses through the control of ABA distribution (Kang et al., 2010; Kuromori et al., 2010), and may also contribute to the coordination between root development and water-saving responses. In addition to ABC transporters, other conserved drought-responsive transport genes included members of the *SWEET*, *NRT1*, and *CWINV* families, involved in carbohydrate and nitrogen transport and metabolism. Their induction under drought conditions suggests a coordinated reprogramming of carbon and nitrogen allocation to sustain root activity and survival. In grapevine, enhanced expression of *SWEET14* and *CWINV* has previously been associated with increased root elongation and improved drought resilience in the 110R rootstock (Yildirim et al., 2018), supporting the idea that resource remobilization constitutes an important component of the conserved root drought response.

Genes associated with ABA metabolism and signaling, cell wall organization, and lipid metabolism also formed a major component of the conserved drought response. Sixteen ABA-related genes involved in ABA biosynthesis, conjugation, signaling, and transport were consistently differentially expressed across the three species, confirming ABA signaling as a central regulator of root drought adaptation. These findings support previous studies demonstrating that root-derived ABA acts as a long-distance signal coordinating whole-plant responses to drought and water deficit (Davies et al., 2005; Abilasha et al., 2021). Beyond its role in stomatal regulation, ABA is widely recognized for modulating root growth, osmotic adjustment, and stress-responsive transcriptional networks that preserve root functionality under dehydration. In grapevine, Cochetel et al. (2020) further demonstrated that roots function not only as drought-responsive tissues but also as active sites of ABA biosynthesis and signaling.

In parallel with the activation of stress-responsive pathways, drought treatment consistently repressed genes associated with defense responses, transcriptional regulation, cytoplasmic translation, and Ca²⁺ signaling in all three species. This coordinated downregulation likely reflects a strategic reallocation of metabolic resources consistent with the stress–growth trade-off framework, whereby plants suppress energetically costly processes to prioritize survival under adverse conditions. The attenuation of biotic defense pathways under abiotic stress has been reported in several grapevine studies (Haider et al., 2017; Hatmi et al., 2018). By contrast, the repression of Ca²⁺ signaling pathways has received comparatively little attention. Dubrovina et al. (2019) previously described altered expression of calmodulin-like (*CML*) genes in *V. amurensis* under abiotic stress conditions. In our study, the downregulation of *CML* and other Ca²⁺ signaling-related genes may indicate that plants had progressed beyond the initial stress perception phase and were actively attenuating signaling activity to avoid unnecessary energy expenditure during sustained drought exposure.

### Species-specific mechanisms: Osmoregulation versus ROS scavenging

*Vitis acerifolia* was the most transcriptionally reactive species in response to water deficit, exhibiting the highest number of DEGs among the studied species. In addition to the conserved drought-responsive pathways, *V. acerifolia* specifically altered the expression of genes involved in solute transport and nutrient uptake, including members of the *NRT1/2*, *AMT2/3*, *GLR*, and *MSL* families. Notably, it was also the only species showing repression of genes associated with effector-triggered immunity (ETI), suggesting a stronger reallocation of resources away from immune functions toward drought tolerance. Together with *V. doaniana*, *V. acerifolia* shared several species-specific DEGs, revealing common adaptive mechanisms. Among these were genes with dual roles in drought response and senescence regulation, such as *RD19* and *SAG12*, which are involved in nutrient remobilization and energy management during stress conditions (Bhat et al., 2019).

Both *V. acerifolia* and *V. doaniana* appeared to invest more strongly in osmoregulatory and hydraulic adjustment mechanisms than *V. candicans*. These two species showed increased expression of genes associated with wax and suberin biosynthesis, including *ESB1*, *ECERIFERUM3*, *Phytosulfokine 3*, and *3-ketoacyl-CoA synthase 1*, as well as several aquaporins (*PIP1-2*, *PIP2-5*, and *TIP*) and MYB transcription factors. Such responses are consistent with the regulation of membrane water transport, suberization, wax deposition, and osmoprotectant accumulation (Gambetta et al., 2012; Yildirim et al., 2018; Gambetta et al., 2020). The stronger induction of these pathways supports the idea that these species preferentially adopt a strategy centered on water conservation and maintenance of root hydraulic function. Although *V. doaniana* displayed physiological responses similar to those of *V. acerifolia*, it was the least transcriptionally reactive species, with the lowest number of DEGs under drought conditions. This intermediate behavior may reflect its proposed origin as a natural hybrid between *V. acerifolia* and *V. candicans* (Zecca et al., 2020).

In contrast, *V. candicans* appeared to rely more strongly on metabolic and oxidative stress protection mechanisms. Although repression of cytoplasmic translation formed part of the conserved drought response, *V. candicans* exhibited a stronger downregulation of genes associated with translational activity, suggesting a more pronounced reduction in energy-consuming metabolic processes. Unlike the other species, enrichment analysis in *V. candicans* did not reveal specific responses related to nutrient uptake or senescence pathways. Instead, this species showed repression of several ABA signaling components, including the ABA receptors *PYR1* and *PYL4*, as well as protein phosphatase inhibitor genes. Given the central role of ABA in stomatal regulation and root-to-shoot drought signaling (Buckley, 2019), these transcriptional differences may partly explain the delayed stomatal closure observed phenotypically in *V. candicans*, particularly since these genes belonged to the green co-expression module significantly correlated with stomatal conductance. Consistent with its stronger metabolic response, *V. candicans* specifically upregulated glycosyltransferase activities, including quercetin 3-*O*-glucosyltransferases, reflecting activation of flavonoid and stilbene biosynthesis pathways involved in oxidative stress mitigation. Corso et al. (2015) previously demonstrated that drought-tolerant grapevine rootstocks exhibit stronger induction of flavonoid biosynthetic genes in roots than drought-sensitive genotypes. Similarly, UDP-glycosyltransferases in *Arabidopsis* have been implicated in abiotic stress adaptation, where their induction correlates with improved drought resistance (Li et al., 2017). It is therefore plausible that *V. candicans* enhances flavonoid glycosylation under drought to modulate antioxidant capacity and limit oxidative damage. In contrast, *V. acerifolia* showed repression of oxidoreductase, *O*-methyltransferase, and peroxidase activities, whereas *V. doaniana* mainly downregulated *O*-methyltransferase-related functions.

These transcriptomic patterns are consistent with the phenotypic and metabolic differences observed among species and collectively support the hypothesis that *V. candicans* relies more heavily on ROS scavenging and secondary metabolism, whereas *V. acerifolia* and *V. doaniana* preferentially invest in osmoregulatory and hydraulic adjustment mechanisms during drought stress.

### Complex gene networks underlie root phenotypes in response to drought

Co-expression network analysis provided additional insights into the molecular regulation underlying drought-responsive phenotypes in grapevine roots. The identified gene modules displayed highly interconnected topologies characterized by numerous hub genes and extensive connectivity among pathways, reflecting the complexity of root phenotypes in response to water deficit. Such network organization highlights the polygenic nature of root traits and emphasizes the challenges associated with identifying robust molecular markers applicable for breeding for drought adaptive phenotypes (de Miguel et al. 2022).

Gene networks associated with shoot-related drought responses were predominantly enriched in signaling and regulatory components, supporting the central role of roots as primary sensors of soil water deficit that coordinate whole-plant acclimation. In particular, the brown module, which correlated with Electron Transport Rate, highlighted the importance of post-transcriptional regulation and organellar RNA editing mediated by pentatricopeptide repeat (PPR) proteins. PPR proteins are increasingly recognized as important regulators of plant stress tolerance through their role in maintaining mitochondrial and chloroplast function under adverse conditions (Wang & Tan, 2025). Several of the PPR genes identified here have previously been associated with drought tolerance in annual species (de Longevialle et al., 2007; Cheng et al., 2023). Among them, *SOAR1* (*Suppressor of the ABAR Overexpressor 1*), a regulator of ABA signaling in *Arabidopsis*, is particularly noteworthy, as its overexpression has been linked to enhanced drought tolerance and improved mitochondrial gene expression under stress conditions (Mei et al., 2014).

More broadly, the co-expression modules associated with both above- and belowground traits reinforced the importance of several core drought-responsive processes identified in the conserved response analysis, notably oxidative stress mitigation, transcriptional regulation, and solute transport. Among the metabolic pathways highlighted, the induction of genes from the trehalose biosynthesis pathway, including *TPS* and *TPP*, underscores the importance of carbohydrate-mediated stress acclimation in grapevine roots. Beyond its role as an osmoprotectant, trehalose functions as a signaling molecule regulating carbon metabolism and ABA-dependent stress responses (Paul et al., 2018; Eh et al., 2024). Overexpression of *TPS* and *TPP* genes has been shown to improve drought tolerance in several species, including *Arabidopsis*, maize, and rice, through membrane stabilization and enhanced antioxidant capacity (Garg et al., 2002; Lin et al., 2019).

Gene modules associated with root phenotypes (RDW, NAR & Ψ) and changing their expression in response to drought highlighted candidate mechanisms involved in root structural and osmotic adjustment. The green-yellow module identified β-galactosidase (*BGAL*) genes as important components of the root drought response. In plants, *BGAL* expression is modulated by abiotic stresses and is primarily associated with cell wall remodeling, while also contributing to osmoregulation (Hou et al., 2021; Rajeev et al., 2024; Buttanri et al., 2025). In parallel, HVA22-like proteins, which are ABA-inducible stress-responsive proteins, were also associated with drought-responsive modules and RDW. These proteins have been implicated in cellular protection and stress resilience under adverse environmental conditions (Zhang et al., 2023). Consistent with our results, Prinsi et al. (2018) previously reported the upregulation of HVA22-like proteins in roots of the grapevine rootstock 101.14 under drought stress.

### Root phenotypes involved in drought adaptation are partly shaped by non-plastic gene expression

Although the transcriptional response to water deficit was largely conserved among the three *Vitis* species, with only subtle species-specific differences in adaptive strategies, variation in root phenotypes associated with drought responses appeared to depend strongly on constitutive gene expression. Our results revealed several significant correlations between drought-responsive root traits and non-plastic patterns of gene expression. In contrast to shoot responses related to gas exchange and photosynthetic activity, which were primarily associated with plastic transcriptional regulation under stress, interspecific differences in root biomass and adventitious root formation under water deficit were more strongly associated with species-dependent constitutive gene expression rather than with drought-induced genes. However, root-osmotic adjustment, a drought adaptation mechanism acting at short-term time scale, was more linked to plastic gene expression than root biomass or the number of adventitious roots that act at longer time-scales (Bernardo et al. 2025).

The random forest analysis further supported this interpretation, as the ten most discriminative genes among the three species were independent of water status and remained stable under drought conditions. Notably, the most discriminative genes were primarily associated with root morphological and physiological phenotypes and they were not differentially expressed under water deficit . These findings highlight the importance of constitutive traits in shaping drought tolerance. From a breeding perspective, these results suggest that constitutively expressed genes associated with root phenotypes can also be considered as targets for the development of drought-resilient grapevine rootstocks.

## Conclusion

Using a comparative multi-omics approach integrating transcriptomic, metabolomic, and physiological analyses, this study investigated root drought responses in three wild *Vitis* species representing different positions along the water use strategy continuum. Our results identified a conserved core drought-responsive program in roots involving solute transport, ABA signaling, oxidative stress mitigation, and metabolic reprogramming, supporting the existence of shared adaptive pathways across *Vitis* species. At the same time, we demonstrated that species-specific responses were associated with distinct transcriptional and metabolic strategies, particularly regarding osmoregulation and antioxidant activity. Importantly, similar physiological outcomes could arise from different molecular responses, highlighting the complexity and multidimensional nature of drought adaptation.

The integration of transcriptomic data with root and shoot phenotypes further revealed that several key drought-related root traits, including root biomass, and adventitious root development, were more strongly associated with constitutive, non-plastic gene expression than with inducible stress-responsive pathways. In particular, the superior root biomass observed in *V. acerifolia* was mostly explained by stable species-dependent expression profiles, while *V. candicans* relied more strongly on inducible antioxidant and secondary metabolism pathways. These findings support our hypothesis that drought adaptation in wild *Vitis* species emerges from the interplay between conserved stress plasticity and intrinsic physiological regulation.

Overall, this work emphasizes the importance of constitutive root traits and species-specific regulatory networks in shaping drought resilience. By moving beyond the traditional comparison of extreme phenotypes, our study provides new insights into the continuum of drought adaptation strategies in grapevine and highlights genes constitutively expressed and associated with root phenotypes as promising targets for the selection and development of drought-resilient rootstocks under future climate scenarios.

## Acknowledgments

We acknowledge Maria Lafargue, Cyril Hevin, Nicolas Hocquard, Jean-Pierre Petit, Laure de Morgadinho Véronique Troadec, Mehmet Koç and Martine Donnart for their help with the plant material and sample preparation. We thank the French INRAE collections from the Grapevine Biological Research Center (Vassal, Montpellier), Ecophysiology and Functional Genomics of Grapevine (EGFV, Bordeaux), and Grapevine Health and Wine Quality (SVQV, Colmar), the Spanish germplasm center “El Encín” and the German collection from the Julius Kühn Institute for kindly providing all the plant material used in this study. We acknowledge the Transcriptome facility (University of Bordeaux, INSERM, PUMA, Neurocentre Magendie, Bordeaux, France) for the advice and technical contribution to the mRNA library preparation, particularly Frédéric Martins. Illumina sequencing was performed at the PGTB (doi:10.15454/1.5572396583599417E12), with the help of Zoé Compagnie and Erwan Guichoux. We are grateful to the genotoul bioinformatics platform Toulouse Occitanie (Bioinfo Genotoul, https://doi.org/10.15454/1.5572369328961167E12) for providing help , computing and storage resources. Finally, we would like to thank Sarah Cookson for sharing her expertise to optimize RNA extraction and her suggestions for the subsequent analysis of sequences.

## Author contribution

E.C: Formal analysis, Writing – original draft. E.R.P: Investigation, Writing – review & editing. JT: Methodology, Writing – review & editing. M. de Miguel: Conceptualization, Funding acquisition, Supervision, Investigation, Writing – review & editing.

## Data availability

Experimental and sequencing data are available upon request

## Supplementary Files

Supplementary Table 1_DifferentialAnalysis

Supplementary Table 2_MapMan _EnrichmentAnalysis

Supplementary Table 3_KEGG_EnrichmentAnalysis

Supplementary Table biorxic4_WGCNA

Supplementary Table 5_RF

Supplementary Table 6_Samples

**Supplementary Figure S1:**
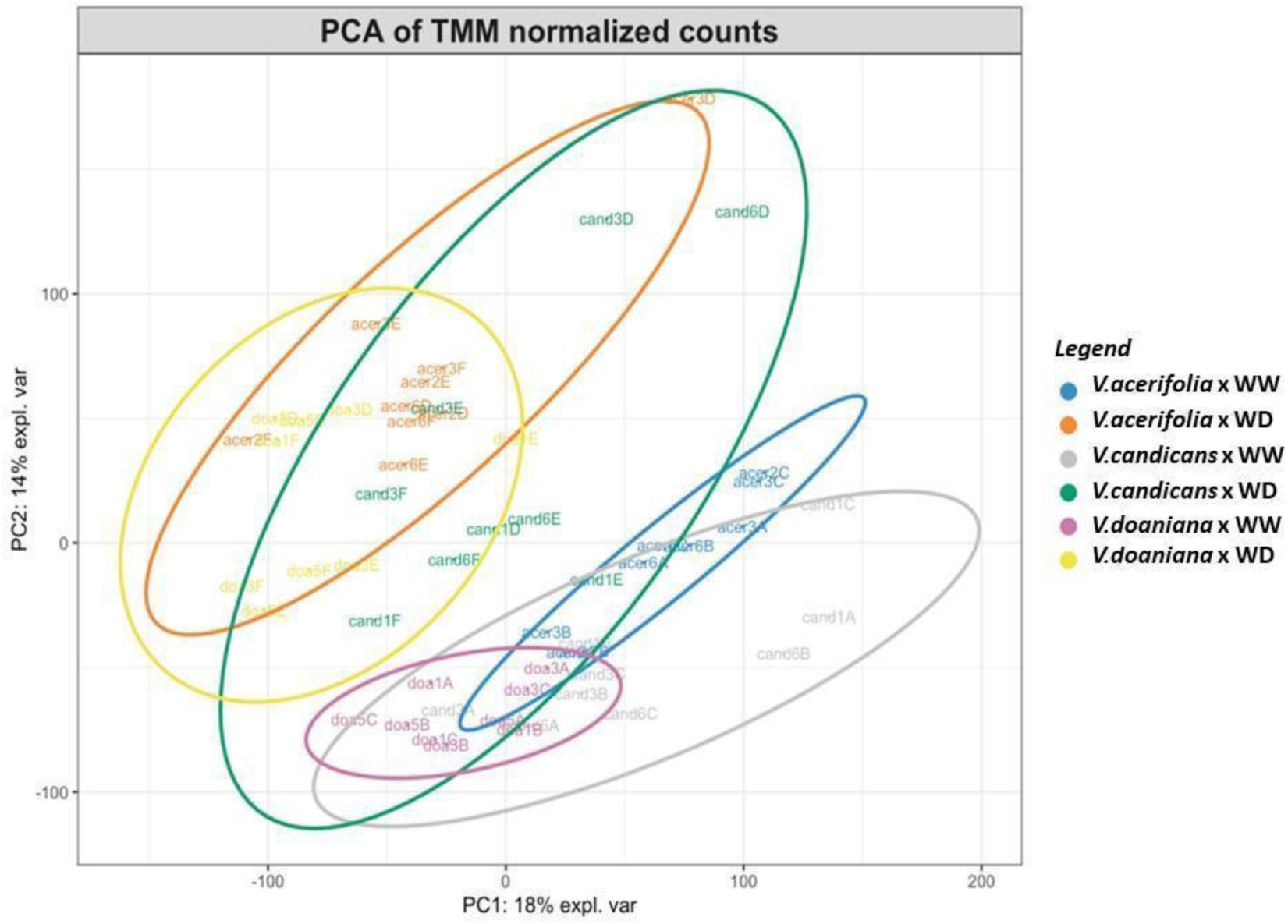
principal component analysis with 21,628 genes expression profiles in three Vitis spp. (V. acerifolia, V. candicans and V. doaniana) comprising three accessions per species, and three replicates per accession and watering treatment.

**Supplementary Figure S2:**
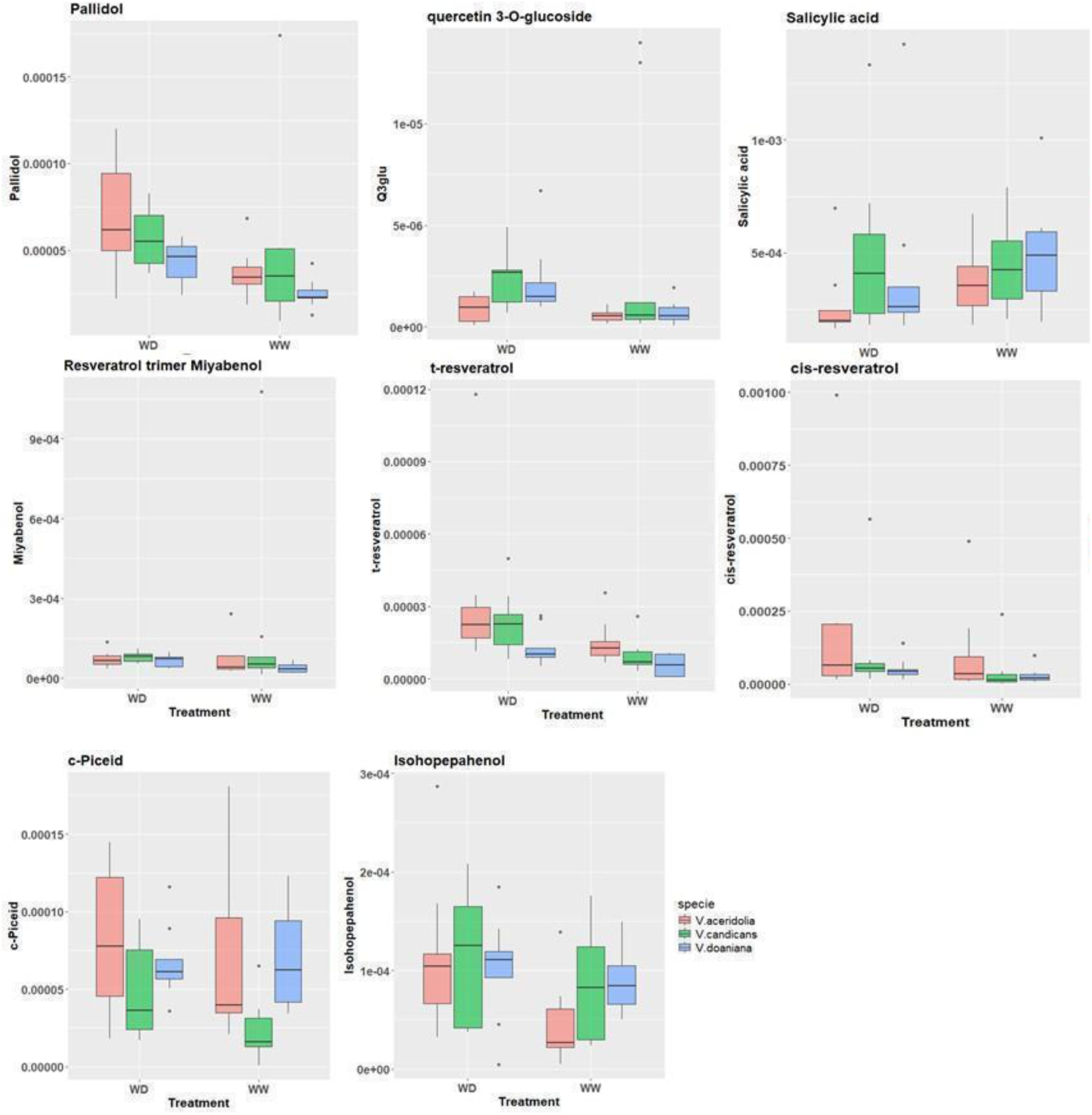
Boxplots of polyphenols with no significant difference between species or treatments

**Supplementary Figure S3:**
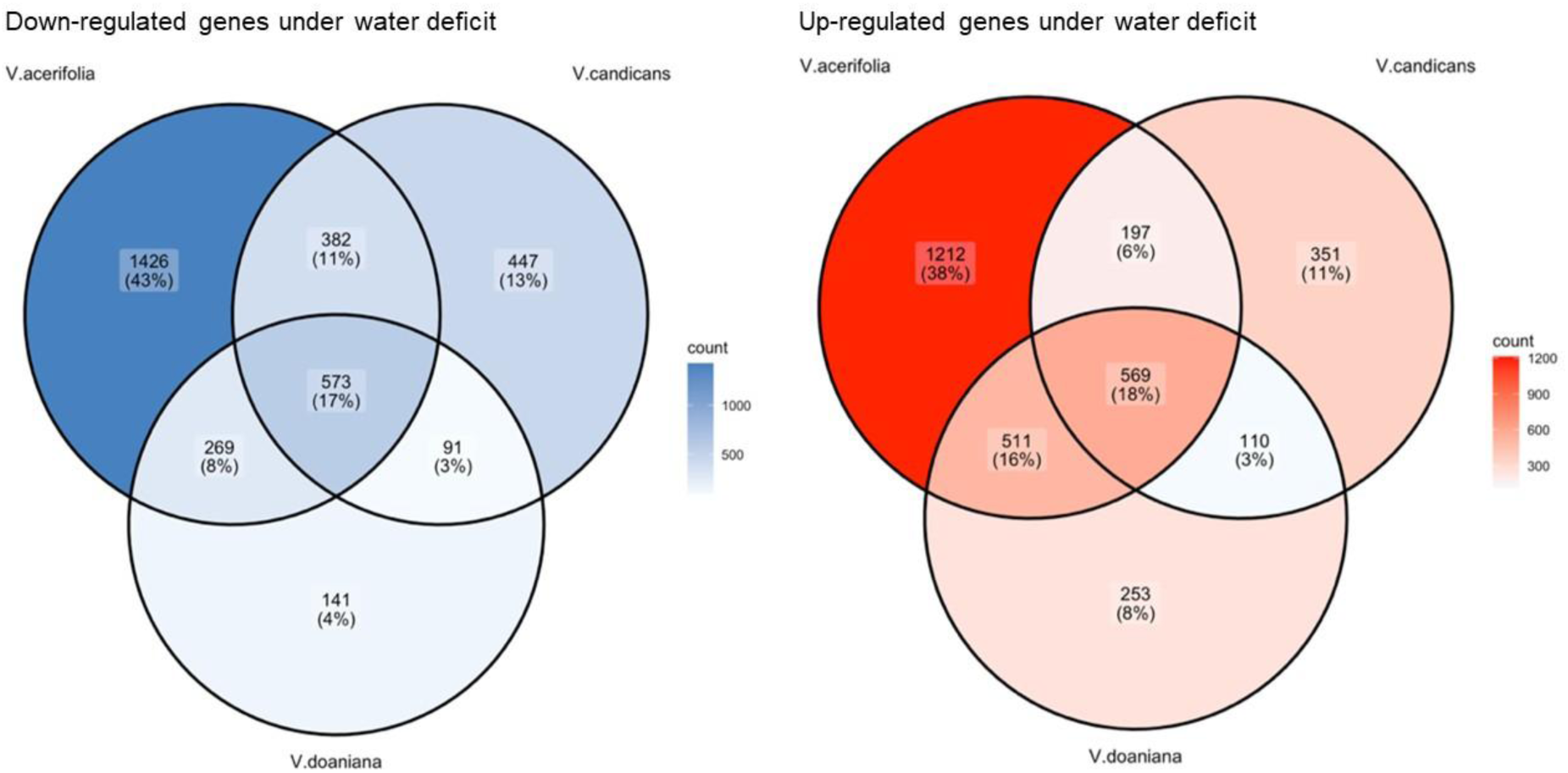
Venn diagrams representing down-regulated genes (left) and up-regulated genes (right) in roots of three different Vitis species under drought conditions

**Supplementary Figure S4:**
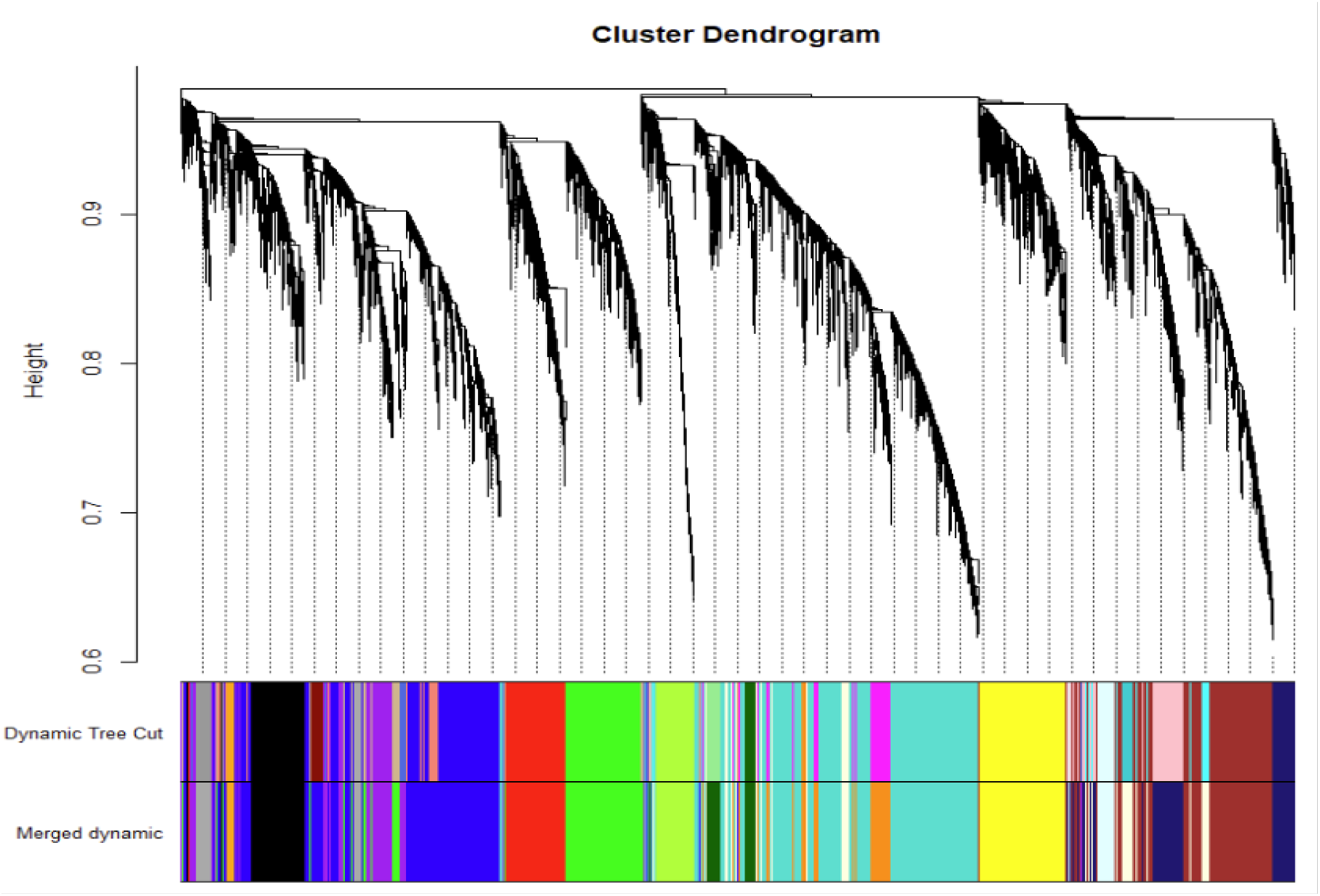
cluster Dendogram from WGCNA analysis

**Supplementary Figure S5:**
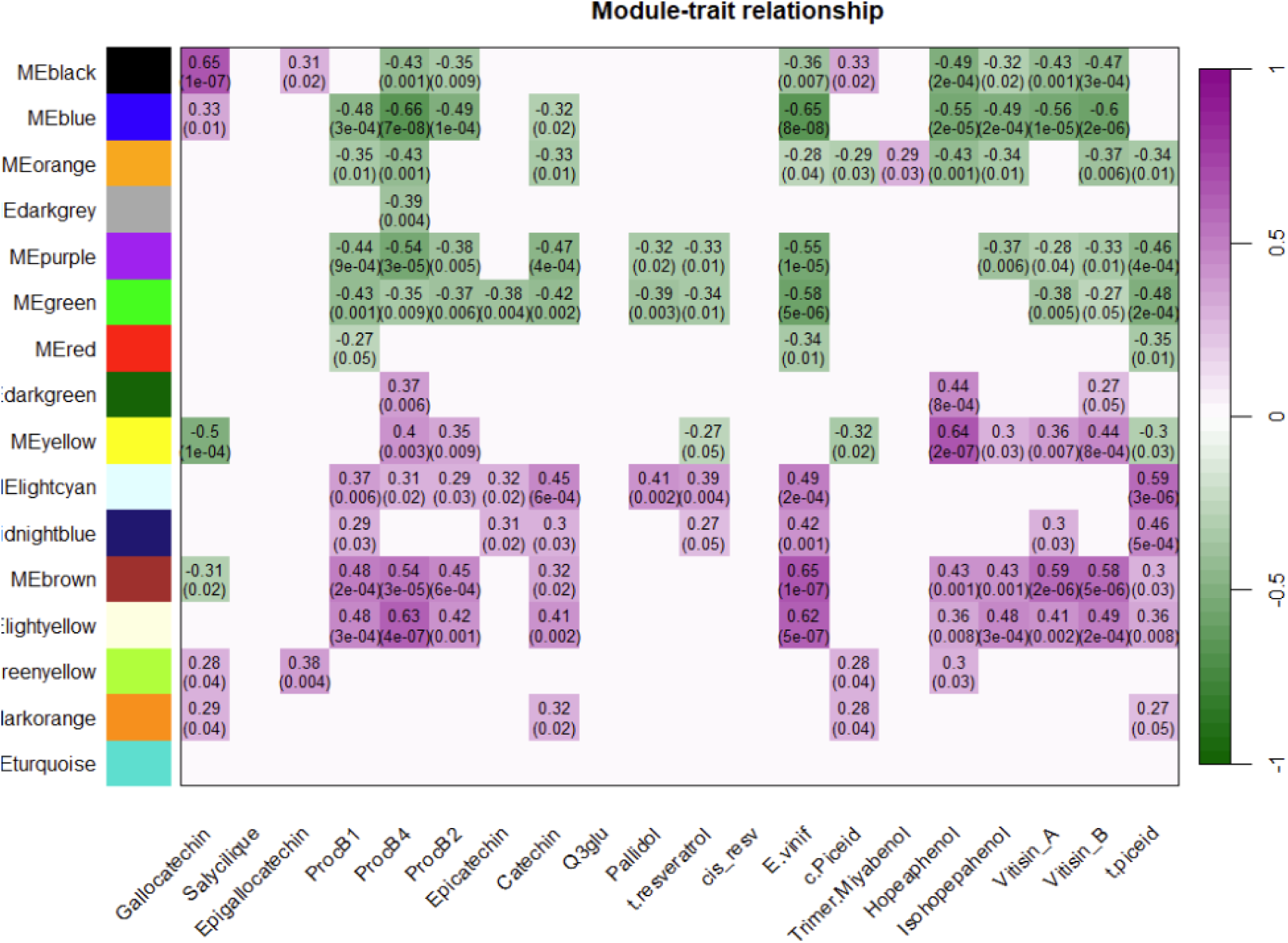
Heatmap for correlation between modules and measured polyphenols in Vitis roots.

**Supplementary Table S1:** Different categories of common differentially expressed genes in roots of three Vitis species under mild drought stress according to Anova-like differential analysis.

|  | Up regulated | Down regulated |
| --- | --- | --- |
| RNA biosynthesis/Transcription regulation | 88 | 156 |
| Enzyme classification - transferases | 106 | 124 |
| Solute transport | 134 | 76 |
| Enzyme classification - oxidoreductases | 89 | 86 |
| Protein modification | 50 | 71 |
| Protein homeostasis | 69 | 50 |
| Phytohormone action | 48 | 41 |
| External stimuli response | 27 | 52 |
| Multi-process regulation | 23 | 43 |
| Enzyme classification - hydrolases | 33 | 32 |
| Cell wall organization | 43 | 19 |
| Nutrient uptake | 24 | 38 |
| Carbohydrate metabolism | 42 | 16 |
| **Lipid metabolism** | 35 | 21 |
| **Photosynthesis** | 19 | 18 |
| **Secondary metabolism** | 20 | 16 |
| **Redox homeostasis** | 13 | 19 |
| **Enzyme classification - lyases** | 13 | 12 |
| **Coenzyme metabolism** | 14 | 8 |
| **Vesicle trafficking** | 10 | 9 |
| **Protein biosynthesis** | 8 | 8 |
| **RNA processing** | 9 | 6 |
| **Amino acid metabolism** | 7 | 4 |
| **Enzyme classification - ligases** | 7 | 3 |
| **Clade-specific metabolism** | 0 | 10 |
| **Plant reproduction** | 5 | 4 |
| **Cell division** | 5 | 2 |
| **Nucleotide metabolism** | 4 | 3 |
| **Enzyme classification - isomerases** | 6 | 0 |
| **Protein translocation** | 0 | 1 |
| **DNA damage response** | 1 | 0 |

## Bibliography

1. Abhilasha, A. and Roy Choudhury, S. (2021) ‘Molecular and physiological perspectives of abscisic acid mediated drought adjustment strategies’, Plants, 10(12), p. 2769. Available at: 10.3390/plants10122769.

2. Alsina, M.M., Smart, D.R., Bauerle, T., De Herralde, F., Biel, C., Stockert, C., Negron, C. and Save, R. (2011) ‘Seasonal changes of whole root system conductance by a drought-tolerant grape root system’, Journal of Experimental Botany, 62(1), pp. 99–109. Available at: 10.1093/jxb/erq247.

3. Andéol Falcon de Longevialle, E.H., Meyer, E.H., Andrés, C., Taylor, N.L., Lurin, C., Millar, A.H. and Small, I.D. (2007) ‘The pentatricopeptide repeat gene OTP43 is required for trans-splicing of the mitochondrial nad1 intron 1 in Arabidopsis thaliana’, The Plant Cell, 19(10), pp. 3256–3265. Available at: 10.1105/tpc.107.054841.

4. Atak, A. (2025) ‘Vitis species for stress tolerance/resistance’, Genetic Resources and Crop Evolution, 72, pp. 2425–2444. Available at: 10.1007/s10722-024-02106-z.

5. Bernardo, S., Marguerit, E., Ollat, N., Gambetta, G.A., Saint Cast, C. and de Miguel, M. (2025) ‘Root system ideotypes: what is the potential for breeding drought-tolerant grapevine rootstocks?’, Journal of Experimental Botany, 76(11), pp. 2970–2984. Available at: 10.1093/jxb/eraf006.

6. Borghi, L., Kang, J., Ko, D., Lee, Y. and Martinoia, E. (2015) ‘The role of ABCG-type ABC transporters in phytohormone transport’, Biochemical Society Transactions, 43(5), pp. 924–930. Available at: 10.1042/BST20150106.

7. Buckley, T.N. (2019) ‘How do stomata respond to water status?’, New Phytologist, 224(1), pp. 21–36.

8. Buttanri, A., Kasapoğlu, A.G., Öner, B.M. et al. (2025) ‘Predicting the role of β-GAL genes in bean under abiotic stress and genome-wide characterization of β-GAL gene family members’, Protoplasma, 262, pp. 365–383. Available at: 10.1007/s00709-024-01998-z.

9. Cantó, C., Menzies, K. J., & Auwerx, J. (2015) ‘NAD(+) Metabolism and the Control of Energy Homeostasis: A Balancing Act between Mitochondria and the Nucleus’, Cell metabolism, 22(1), pp. 31–53. 10.1016/j.cmet.2015.05.023

10. Cantu, D., Massonnet, M. and Cochetel, N. (2024) ‘The wild side of grape genomics’, Trends in Genetics, 40, pp. 601–612.

11. Carvalho, L.C., Coito, J.L., Gonçalves, E.F., Chaves, M.M. and Amâncio, S. (2016) ‘Differential physiological response of the grapevine varieties Touriga Nacional and Trincadeira to combined heat, drought and light stresses’, Plant Biology, 18, pp. 101–111.

12. Chaudhry, S. and Sidhu, G.P.S. (2022) ‘Climate change regulated abiotic stress mechanisms in plants: a comprehensive review’, Plant Cell Reports, 41(1), pp. 1–31.

13. Chaves, M. and Davies, B. (2010) ‘Drought effects and water use efficiency: improving crop production in dry environments’, Functional Plant Biology, 37(2), pp. iii–vi.

14. Cheng, L., Lu, K. and Wang, C. (2023) ‘Research progress of PPR protein in plant abiotic stress response’, Scientia Agricultura Sinica, 56(24), pp. 4801–4813. Available at: 10.3864/j.issn.0578-1752.2023.24.001.

15. Christmann, A., Weiler, E.W., Steudle, E. and Grill, E. (2007) ‘A hydraulic signal in root-to-shoot signalling of water shortage’, The Plant Journal, 52(1), pp. 167–174.

16. Climate Change 2023: Synthesis Report. Contribution of Working Groups I, II and III to the Sixth Assessment Report of the Intergovernmental Panel on Climate Change., IPCC, Geneva, Switzerland, pp. 35–115

17. Cochetel, N., Ghan, R., Toups, H.S., Degu, A., Tillett, R.L., Schlauch, K.A. and Cramer, G.R. (2020) ‘Drought tolerance of the grapevine, Vitis champinii cv. Ramsey, is associated with higher photosynthesis and greater transcriptomic responsiveness of abscisic acid biosynthesis and signaling’, BMC Plant Biology, 20(1), p. 55.

18. Comas, L.H., Becker, S.R., Cruz, V.M.V., Byrne, P.F. and Dierig, D.A. (2013) ‘Root traits contributing to plant productivity under drought’, Frontiers in Plant Science, 4, p. 442.

19. Condon, A.G., Richards, R.A., Rebetzke, G.J. and Farquhar, G.D. (2004) ‘Breeding for high water-use efficiency’, Journal of Experimental Botany, 55(407), pp. 2447–2460.

20. Cookson, S.J. and Ollat, N. (2013) ‘Grafting with rootstocks induces extensive transcriptional re-programming in the shoot apical meristem of grapevine’, BMC Plant Biology, 13, p. 147. Available at: 10.1186/1471-2229-13-147.

21. Coupel-Ledru, A., Tyerman, S.D., Masclef, D., Lebon, E., Christophe, A., Edwards, E.J. and Simonneau, T. (2017) ‘Abscisic acid down-regulates hydraulic conductance of grapevine leaves in isohydric genotypes only’, Plant Physiology, 175(3), pp. 1121–1134. Available at: 10.1104/pp.17.00698.

22. Corso, M., Vannozzi, A., Maza, E., Vitulo, N., Meggio, F., Pitacco, A., Telatin, A., D’Angelo, M., Feltrin, E., Negri, A.S., Prinsi, B., Valle, G., Ramina, A., Bouzayen, M., Bonghi, C. and Lucchin, M. (2015) ‘Comprehensive transcript profiling of two grapevine rootstock genotypes contrasting in drought susceptibility links the phenylpropanoid pathway to enhanced tolerance’, Journal of Experimental Botany, 66(19), pp. 5739–5752. Available at: 10.1093/jxb/erv274.

23. Davies, W.J., Kudoyarova, G. and Hartung, W. (2005) ‘Long-distance ABA signaling and its relation to other signaling pathways in the detection of soil drying and the mediation of the plant’s response to drought’, Journal of Plant Growth Regulation, 24(4), pp. 285–295. Available at: 10.1007/s00344-005-0103-1.

24. Dayer, S., Herrera, J.C., Dai, Z., Burlett, R., Lamarque, L.J., Delzon, S., Bortolami, G., Cochard, H. and Gambetta, G.A. (2020) ‘The sequence and thresholds of leaf hydraulic traits underlying grapevine varietal differences in drought tolerance’, Journal of Experimental Botany, 71(14), pp. 4333–4344. Available at: 10.1093/jxb/eraa186.

25. Degu, A., Hochberg, U., Wong, D.C.J. et al. (2019) ‘Swift metabolite changes and leaf shedding are milestones in the acclimation process of grapevine under prolonged water stress’, BMC Plant Biology, 19, p. 69. Available at: 10.1186/s12870-019-1652-y.

26. Delzon, S. (2015) ‘New insight into leaf drought tolerance’, Functional Ecology, 29(10), pp. 1247–1249.

27. de Miguel, M., Rodríguez-Quilón, I., Heuertz, M., Hurel, A., Grivet, D., Jaramillo-Correa, J.P., Vendramin, G.G., Plomion, C., Majada, J., Alía, R., Eckert, A.J. and González-Martínez, S.C. (2022) ‘Polygenic adaptation and negative selection across traits, years and environments in a long-lived plant species (Pinus pinaster Ait., Pinaceae)’, Molecular Ecology, 31, pp. 2089–2105. Available at: 10.1111/mec.16367.

28. Dubrovina, A.S., Aleynova, O.A., Ogneva, Z.V., Suprun, A.R., Ananev, A.A. and Kiselev, K.V. (2019) ‘The effect of abiotic stress conditions on expression of calmodulin (CaM) and calmodulin-like (CML) genes in wild-growing grapevine Vitis amurensis’, Plants, 8(12), p. 602. Available at: 10.3390/plants8120602.

29. Eh, T.J., Lei, P., Phyon, J.M., Kim, H.I., Xiao, Y., Ma, L., Li, J., Bai, Y., Ji, X., Jin, G. and Meng, F. (2024) ‘The AaERF64-AaTPPA module participates in cold acclimatization of *Actinidia arguta* (Sieb. et Zucc.) Planch ex Miq.’, Molecular Breeding, 44(6), p. 43. Available at: 10.1007/s11032-024-01475-8.

30. Flamini R, Mattivi F, De Rosso M, Arapitsas P, Bavaresco L. (2013). ‘Advanced knowledge of three important classes of grape phenolics: anthocyanins, stilbenes and flavonols’. Int J Mol Sci. 14(10), p. 19651–69. doi: 10.3390/ijms141019651.

31. Flor, L., Toro, G., Carriquí, M., Buesa, I., Sabater, A., Medrano, H. and Escalona, J.M. (2025) ‘Impact of severe water stress on drought resistance mechanisms and hydraulic vulnerability segmentation in grapevine: the role of rootstock’, Journal of Experimental Botany, 76(11), pp. 3141–3157. Available at: 10.1093/jxb/eraf044.

32. Gambetta, G.A., Fei, J., Rost, T.L., Knipfer, T., Matthews, M.A., Shackel, K.A., Walker, M.A. and McElrone, A.J. (2013) ‘Water uptake along the length of grapevine fine roots: developmental anatomy, tissue-specific aquaporin expression, and pathways of water transport’, Plant Physiology, 163(3), pp. 1254–1265. Available at: 10.1104/pp.113.221283.

33. Gambetta, G.A., Knipfer, T., Fricke, W. and McElrone, A.J. (2017) ‘Aquaporins and root water uptake’, in Plant Aquaporins: From Transport to Signaling. Cham: Springer International Publishing, pp. 133–153.

34. Gambetta, G.A., Herrera, J.C., Dayer, S., Feng, Q., Hochberg, U. and Castellarin, S.D. (2020) ‘The physiology of drought stress in grapevine: towards an integrative definition of drought tolerance’, Journal of Experimental Botany, 71(16), pp. 4658–4676. Available at: 10.1093/jxb/eraa245.

35. Garg, A.K., Kim, J.K., Owens, T.G., Ranwala, A.P., Choi, Y.D., Kochian, L.V. and Wu, R.J. (2002) ‘Trehalose accumulation in rice plants confers high tolerance levels to different abiotic stresses’, Proceedings of the National Academy of Sciences, 99(25), pp. 15898–15903. Available at: 10.1073/pnas.252637799.

36. Gargallo-Garriga, A., Sardans, J., Pérez-Trujillo, M. (2014) ‘Opposite metabolic responses of shoots and roots to drought’, Sci Rep, 4, p. 6829. 10.1038/srep06829

37. Gautier, A.T., Cochetel, N., Merlin, I. et al. (2020) ‘Scion genotypes exert long distance control over rootstock transcriptome responses to low phosphate in grafted grapevine’, BMC Plant Biology, 20, p. 367. Available at: 10.1186/s12870-020-02578-y.

38. Gautier, A.T., Merlin, I., Doumas, P., Cochetel, N., Mollier, A., Vivin, P., Lauvergeat, V., Péret, B. and Cookson, S.J. (2021) ‘Identifying roles of the scion and the rootstock in regulating plant development and functioning under different phosphorus supplies in grapevine’, Environmental and Experimental Botany, 185, p. 104405. Available at: 10.1016/j.envexpbot.2021.104405.

39. Ge, S.X., Jung, D. and Yao, R. (2020) ‘ShinyGO: a graphical gene-set enrichment tool for animals and plants’, Bioinformatics, 36(8), pp. 2628–2629. Available at: 10.1093/bioinformatics/btz931.

40. Goodger, J.Q. and Schachtman, D.P. (2010) ‘Re-examining the role of ABA as the primary long-distance signal produced by water-stressed roots’, Plant Signaling & Behavior, 5(10), pp. 1298–1301.

41. Gregory, J.P. (1974) ‘Vegetation classification by reference to strategies’, Nature, 250, pp. 26–31.

42. Gupta, A., Rico-Medina, A. and Caño-Delgado, A.I. (2020) ‘The physiology of plant responses to drought’, Science, 368(6488), pp. 266–269.

43. Haider, M.S., Zhang, C., Kurjogi, M.M., Pervaiz, T., Zheng, T., Zhang, C., Lide, C., Shangguan, L. and Fang, J. (2017) ‘Insights into grapevine defense response against drought as revealed by biochemical, physiological and RNA-Seq analysis’, Scientific Reports, 7(1), p. 13134. Available at: 10.1038/s41598-017-13464-4.

44. Hanzouli, F., Daldoul, S., Zemni, H., Boubakri, H., Vincenzi, S., Mliki, A. and Gargouri, M. (2025) ‘Stilbene production as part of drought adaptation mechanisms in cultivated grapevine (*Vitis vinifera* L.) roots modulates antioxidant status’, Plant Biology, 27(1), pp. 102–115.

45. Hatmi, S., Villaume, S., Trotel-Aziz, P., Barka, E.A., Clément, C. and Aziz, A. (2018) ‘Osmotic stress and ABA affect immune response and susceptibility of grapevine berries to gray mold by priming polyamine accumulation’, Frontiers in Plant Science, 9, p. 1010. Available at: 10.3389/fpls.2018.01010.

46. He, X., Xu, L., Pan, C., Gong, C., Wang, Y., Liu, X. and Yu, Y. (2020) ‘Drought resistance of *Camellia oleifera* under drought stress: changes in physiology and growth characteristics’, PLoS ONE, 15(7), p. e0235795.

47. Hochberg, U., Degu, A., Toubiana, D., Gendler, T., Nikoloski, Z., Rachmilevitch, S. and Fait, A. (2013) ‘Metabolite profiling and network analysis reveal coordinated changes in grapevine water stress response’, BMC Plant Biology, 13(1), p. 184. Available at: 10.1186/1471-2229-13-184.

48. Hou, F., Du, T., Qin, Z. et al. (2021) ‘Genome-wide in silico identification and expression analysis of beta-galactosidase family members in sweetpotato (*Ipomoea batatas* (L.) Lam)’, BMC Genomics, 22, p. 140. Available at: 10.1186/s12864-021-07436-1.

49. Kamiya, T., Borghi, M., Wang, P., Danku, J.M.C., Kalmbach, L., Hosmani, P.S., Naseer, S., Fujiwara, T., Geldner, N. and Salt, D.E. (2015) ‘The MYB36 transcription factor orchestrates Casparian strip formation’, Proceedings of the National Academy of Sciences of the United States of America, 112(33), pp. 10533–10538. Available at: 10.1073/pnas.1507691112.

50. Kanehisa, M., Araki, M., Goto, S., Hattori, M., Hirakawa, M., Itoh, M., Katayama, T., Kawashima, S., Okuda, S., Tokimatsu, T. and Yamanishi, Y. (2007) ‘KEGG for linking genomes to life and the environment’, Nucleic Acids Research, 36(suppl_1), pp. D480–D484. Available at: 10.1093/nar/gkm882.

51. Kang, J., Hwang, J.U., Lee, M., Kim, Y.Y., Assmann, S.M., Martinoia, E. and Lee, Y. (2010) ‘PDR-type ABC transporter mediates cellular uptake of the phytohormone abscisic acid’, Proceedings of the National Academy of Sciences, 107(5), pp. 2355–2360. Available at: 10.1073/pnas.0909222107.

52. Khadka, V.S., Vaughn, K., Xie, J., Swaminathan, P., Ma, Q., Cramer, G.R. and Fennell, A.Y. (2019) ‘Transcriptomic response is more sensitive to water deficit in shoots than roots of *Vitis riparia* (Michx.)’, BMC Plant Biology, 19(1), p. 72.

53. Konecny, T., Asatryan, A. and Binder, H. (2025) ‘Responding to stress: diversity and resilience of grapevine in a changing climate under the perspective of omics research’, International Journal of Molecular Sciences, 26(16), p. 7877.

54. Kou, X., Zhao, Z., Xu, X., Li, C., Wu, J. and Zhang, S. (2024) ‘Identification and expression analysis of ATP-binding cassette (ABC) transporters revealed its role in regulating stress response in pear (*Pyrus bretchneideri*)’, BMC Genomics, 25(1), p. 169. Available at: 10.1186/s12864-024-10063-1.

55. Kuromori, T., Miyaji, T., Yabuuchi, H., Shimizu, H., Sugimoto, E., Kamiya, A., Moriyama, Y. and Shinozaki, K. (2010) ‘ABC transporter AtABCG25 is involved in abscisic acid transport and responses’, Proceedings of the National Academy of Sciences, 107(5), pp. 2361–2366. Available at: 10.1073/pnas.0912516107.

56. Lesk, C., Rowhani, P. and Ramankutty, N. (2016) ‘Influence of extreme weather disasters on global crop production’, Nature, 529, pp. 84–87. Available at: 10.1038/nature16467.

57. Li, P., Li, Y.J., Zhang, F.J., Zhang, G.Z., Jiang, X.Y., Yu, H.M. and Hou, B.K. (2017) ‘The Arabidopsis UDP-glycosyltransferases UGT79B2 and UGT79B3 contribute to cold, salt and drought stress tolerance via modulating anthocyanin accumulation’, The Plant Journal, 89(1), pp. 85–103. Available at: 10.1111/tpj.13324.

58. Lin, Q., Yang, J., Wang, Q., Zhu, H., Chen, Z., Dao, Y. and Wang, K. (2019) ‘Overexpression of the trehalose-6-phosphate phosphatase family gene AtTPPF improves the drought tolerance of Arabidopsis thaliana’, BMC Plant Biology, 19(1), p. 381. Available at: 10.1186/s12870-019-1986-5.

59. Lin, Y., Liu, S., Fang, X., Ren, Y., You, Z., Xia, J., Hakeem, A., Yang, Y., Wang, L., Fang, J. and Shangguan, L. (2023) ‘The physiology of drought stress in two grapevine cultivars: photosynthesis, antioxidant system, and osmotic regulation responses’, Physiologia Plantarum, 175(5), p. e14005.

60. Liu W, Feng Y, Yu S, Fan Z, Li X, Li J, Yin H., (2021), ‘The Flavonoid Biosynthesis Network in Plants’. Int J Mol Sci. (26)22, p.12824. doi: 10.3390/ijms222312824.

61. Loupit, G., Valls Fonayet, J., Tran, J., Garcia, V., Hummel, I., Petriacq, P., Gallusci, P., Berger, M., Franc, C., de Revel, G., Ollat, N. and Cookson, S.J. (2025) ‘Graft union formation involves interactions among bud signals, carbon availability, dormancy release, wound responses and non-self-communication in grapevine’, The Plant Journal, 122, p. e70244. Available at: 10.1111/tpj.70244.

62. Ma, D. and Constabel, C.P. (2019) ‘MYB repressors as regulators of phenylpropanoid metabolism in plants’, Trends in Plant Science, 24(3), pp. 275–289. Available at: 10.1016/j.tplants.2018.12.003.

63. Malcheska, F., Ahmad, A., Batool, S., Müller-Moulé, J., Kreuzwieser, J., Randewig, D., Hänsch, R., Mendel, R.R., Hell, R. and Wirtz, M. (2017) ‘Drought-enhanced xylem sap sulfate closes stomata by affecting ALMT12 and guard cell ABA synthesis’, Plant Physiology, 174(2), pp. 798–814.

64. Martorell, S., Díaz-Espejo, A., Tomàs, M., Pou, A., El Aou-ouad, H., Escalona, J.M., Vadell, J., Ribas-Carbó, M., Flexas, J. and Medrano, H. (2015) ‘Differences in water-use-efficiency between two *Vitis vinifera* cultivars (Grenache and Tempranillo) explained by the combined response of stomata to hydraulic and chemical signals during water stress’, Agricultural Water Management, 156, pp. 1–9. Available at: 10.1016/j.agwat.2015.03.011.

65. Mei, C., Jiang, S.C., Lu, Y.F., Wu, F.Q., Yu, Y.T., Liang, S., Feng, X.J., Portoles Comeras, S., Lu, K., Wu, Z., Wang, X.F. and Zhang, D.P. (2014) ‘Arabidopsis pentatricopeptide repeat protein SOAR1 plays a critical role in abscisic acid signalling’, Journal of Experimental Botany, 65(18), pp. 5317–5330. Available at: 10.1093/jxb/eru293.

66. Monteiro-Batista, R. D. C., Siqueira, J. A., da Fonseca-Pereira, P., Barreto, P., Feitosa-Araujo, E., Araújo, W. L., & Nunes-Nesi, A. (2025). ‘Potential roles of mitochondrial carrier proteins in plant responses to abiotic stress’, Journal of Experimental Botany, 76(17), 4760–4770. 10.1093/jxb/eraf032.

67. Morales-Cruz, A., Aguirre-Liguori, J.A., Zhou, Y. et al. (2021) ‘Introgression among North American wild grapes (*Vitis*) fuels biotic and abiotic adaptation’, Genome Biology, 22, p. 254. Available at: 10.1186/s13059-021-02467-z.

68. Nio, S.A., Ludong, D.P.M. and Wade, L.J. (2018) ‘Comparison of leaf osmotic adjustment expression in wheat (*Triticum aestivum* L.) under water deficit between the whole plant and tissue levels’, Agriculture and Natural Resources, 52(1), pp. 33–38. Available at: 10.1016/j.anres.2018.03.003.

69. Osakabe, Y., Osakabe, K., Shinozaki, K. and Tran, L.S.P. (2014) ‘Response of plants to water stress’, Frontiers in Plant Science, 5, p. 86.

70. Patin, E.R., Pérez-López, U., del Sol Iturralde, A., Valls-Fonayet, J., Pétriacq, P., Gastou, P., Tandonnet, J.P., Larrey, M., Vivin, P., Marguerit, E., Ollat, N. and de Miguel, M. (2025) ‘Linking the genetic diversity of root traits and drought responses in wild *Vitis* species’, Plant Stress, p. 100964.

71. Paul, M.J., Gonzalez-Uriarte, A., Griffiths, C.A. and Hassani-Pak, K. (2018) ‘The role of trehalose 6-phosphate in crop yield and resilience’, Plant Physiology, 177(1), pp. 12–23. Available at: 10.1104/pp.17.01634.

72. Prinsi, B., Negri, A.S., Failla, O. et al. (2018) ‘Root proteomic and metabolic analyses reveal specific responses to drought stress in differently tolerant grapevine rootstocks’, BMC Plant Biology, 18, p. 126. Available at: 10.1186/s12870-018-1343-0.

73. Rajeev, A.C.G.P. and Pan, A. (2024) ‘Integrated gene expression and network analysis identify drought-response genes and pathways in *Solanum*: a computational study’, Discover Applied Sciences, 6, p. 468. Available at: 10.1007/s42452-024-06025-7.

74. Riaz, S., Pap, D., Uretsky, J., Laucou, V., Boursiquot, J. M., Kocsis, L., & Andrew Walker, M. (2019). ‘Genetic diversity and parentage analysis of grape rootstocks’, Theoretical and Applied Genetics, 132(6), 1847–1860. 10.1007/s00122-019-03320-5

75. Rossdeutsch, L., Edwards, E., Cookson, S.J., Barrieu, F., Gambetta, G.A., Delrot, S. and Ollat, N. (2016) ‘ABA-mediated responses to water deficit separate grapevine genotypes by their genetic background’, BMC Plant Biology, 16(1), p. 91.

76. Sadok, W., Naudin, P., Boussuge, B., Muller, B., Welcker, C. and Tardieu, F. (2007) ‘Leaf growth rate per unit thermal time follows QTL-dependent daily patterns in hundreds of maize lines under naturally fluctuating conditions’, Plant, Cell & Environment, 30, pp. 135–146. Available at: 10.1111/j.1365-3040.2006.01611.x.

77. Schultz, H.R. (2003) ‘Differences in hydraulic architecture account for near-isohydric and anisohydric behaviour of two field-grown *Vitis vinifera* L. cultivars during drought’, *Plant*, Cell & Environment, 26(8), pp. 1393–1405. Available at: 10.1046/j.1365-3040.2003.01064.x.

78. Shelden, M.C., Howitt, S.M., Kaiser, B.N. and Tyerman, S.D. (2009) ‘Identification and functional characterisation of aquaporins in the grapevine, *Vitis vinifera*’, Functional Plant Biology, 36(12), pp. 1065–1078.

79. Shinozaki, K. and Yamaguchi-Shinozaki, K. (2007) ‘Gene networks involved in drought stress response and tolerance’, Journal of Experimental Botany, 58(2), pp. 221–227. Available at: 10.1093/jxb/erl164.

80. Tardieu, F., Simonneau, T. and Muller, B. (2018) ‘The physiological basis of drought tolerance in crop plants: a scenario-dependent probabilistic approach’, Annual Review of Plant Biology, 69, pp. 733–759.

81. Tombesi, S., Nardini, A., Frioni, T. et al. (2015) ‘Stomatal closure is induced by hydraulic signals and maintained by ABA in drought-stressed grapevine’, Scientific Reports, 5, p. 12449. Available at: 10.1038/srep12449.

82. Uga, Y., Sugimoto, K., Ogawa, S., Rane, J., Ishitani, M., Hara, N., Kitomi, Y., Inukai, Y., Ono, K., Kanno, N., Inoue, H., Takehisa, H., Motoyama, R., Nagamura, Y., Wu, J., Matsumoto, T., Takai, T., Okuno, K. and Yano, M. (2013) ‘Control of root system architecture by DEEPER ROOTING 1 increases rice yield under drought conditions’, Nature Genetics, 45(9), pp. 1097–1102. Available at: 10.1038/ng.2725.

83. Uga, Y., Sugimoto, K., Ogawa, S., Rane, J., Ishitani, M., Hara, N., Kitomi, Y., Inukai, Y., Ono, K., Kanno, N., Inoue, H., Takehisa, H., Motoyama, R., Nagamura, Y., Wu, J., Matsumoto, T., Takai, T., Okuno, K. and Yano, M. (2013) ‘Control of root system architecture by DEEPER ROOTING 1 increases rice yield under drought conditions’, Nature Genetics, 45(9), pp. 1097–1102. Available at: 10.1038/ng.2725.

84. Van Leeuwen, C., Destrac-Irvine, A., Dubernet, M., Duchêne, E., Gowdy, M., Marguerit, E., Pieri, P., Parker, A., De Resseguier, L. and Ollat, N. (2019) ‘An update on the impact of climate change in viticulture and potential adaptations’, Agronomy, 9(9), p. 514.

85. Vannozzi A, Dry IB, Fasoli M, Zenoni S, Lucchin M. (2012) ‘Genome-wide analysis of the grapevine stilbene synthase multigenic family: genomic organization and expression profiles upon biotic and abiotic stresses’. BMC Plant Biol., 3(12), p. 130. doi: 10.1186/1471-2229-12-130.

86. Velt, A., Frommer, B., Blanc, S., Holtgräwe, D., Duchêne, E., Dumas, V., Grimplet, J., Hugueney, P., Kim, C., Lahaye, M., Matus, J.T., Navarro-Payá, D., Orduña, L., Tello-Ruiz, M.K., Vitulo, N., Ware, D. and Rustenholz, C. (2023) ‘An improved reference of the grapevine genome reasserts the origin of the PN40024 highly homozygous genotype’, G3: Genes, Genomes, Genetics, 13(5), jkad067. Available at: 10.1093/g3journal/jkad067.

87. Walker, M.A., Heinitz, C., Riaz, S. and Uretsky, J. (2019) ‘Grape taxonomy and germplasm’, in The Grapevine Genome. Cham: Springer International Publishing, pp. 25–38.

88. Wang, Y. and Tan, B.C. (2025) ‘Pentatricopeptide repeat proteins in plants: cellular functions, action mechanisms, and potential applications’, Plant Communications, 6(2). Available at: 10.1016/j.xplc.2024.101203.

89. Wang, X.C., Wu, J., Guan, M.L., Zhao, C.H., Geng, P. and Zhao, Q. (2020) ‘Arabidopsis MYB4 plays dual roles in flavonoid biosynthesis’, The Plant Journal, 101(3), pp. 637–652. Available at: 10.1111/tpj.14570.

90. Yadav, S., Kalwan, G., Gill, S.S. and Jain, P.K. (2025) ‘The ABC transporters and their epigenetic regulation under drought stress in chickpea’, Plant Physiology and Biochemistry, 223, p. 109903. Available at: 10.1016/j.plaphy.2025.109903.

91. Yang, X., Cui, X., Chang, J., Wang, J., Wang, Y., Liu, H., Wang, Y., Chen, Y., Yang, Y., Yao, D. et al. (2024) ‘Variations in protein and gene expression involved in the pathways of carbohydrate, abscisic acid, and ATP-binding cassette transporter in soybean roots under drought stress’, Agronomy, 14(4), p. 843. Available at: 10.3390/agronomy14040843.

92. Yıldırım, K., Yağcı, A., Sucu, S. and Tunç, S. (2018) ‘Responses of grapevine rootstocks to drought through altered root system architecture and root transcriptomic regulations’, Plant Physiology and Biochemistry, 127, pp. 256–268. Available at: 10.1016/j.plaphy.2018.03.034.

93. Zecca, G., Grassi, F., Tabidze, V., Pipia, I., Kotorashvili, A., Kotaria, N. and Beridze, T. (2020) ‘Dates and rates in grape’s plastomes: evolution in slow motion’, Current Genetics, 66(1), pp. 123–140. Available at: 10.1007/s00294-019-01004-7.

94. Zhang, L., Xiao, J., Zhou, Y., Zheng, Y., Li, J. and Xiao, H. (2016) ‘Drought events and their effects on vegetation productivity in China’, Ecosphere, 7(12), p. e01591. Available at: 10.1002/ecs2.1591.

95. Zhang, H., Yuan, Y., Xing, H., Xin, M., Saeed, M., Wu, Q., Wu, J., Zhuang, T., Zhang, X., Mao, L., Sun, X., Song, X. and Wang, Z. (2023) ‘Genome-wide identification and expression analysis of the HVA22 gene family in cotton and functional analysis of GhHVA22E1D in drought and salt tolerance’, Frontiers in Plant Science, 14, p. 1139526. Available at: 10.3389/fpls.2023.1139526.

